# Exact distribution of the quantal content in synaptic transmission

**DOI:** 10.1101/2022.12.28.522121

**Authors:** Krishna Rijal, Nicolas I.C. Müller, Eckhard Friauf, Abhyudai Singh, Ashok Prasad, Dibyendu Das

**Affiliations:** Department of Physics, Indian Institute of Technology Bombay, Powai, Mumbai 400076, India; Animal Physiology Group, Department of Biology, University of Kaiserslautern, Kaiserslautern, Germany; Departments of Electrical and Computer Engineering, Biomedical Engineering and Mathematical Sciences, University of Delaware, Newark, DE 19716, USA; Department of Chemical and Biological Engineering, Colorado State University, Fort Collins, Colorado, USA

## Abstract

During electro-chemical signal transmission through synapses, triggered by an action potential (AP), a stochastic number of synaptic vesicles (SV), called the *quantal content*, release neurotransmitters in the synaptic cleft. It is widely accepted that the quantal content probability distribution is a binomial based on the number of ready-release SVs in the pre-synaptic terminal. But the latter number itself fluctuates due to its stochastic replenishment, hence the actual distribution of quantal content is unknown. We show that exact distribution of quantal content can be derived for general stochastic AP inputs in the steady-state. For fixed interval AP train, we prove that the distribution is a binomial, and corroborate our predictions by comparison with electrophysiological recordings from MNTB-LSO synapses of juvenile mice. For a Poisson train, we show that the distribution is non-binomial. Moreover, we find exact moments of the quantal content in the Poisson and other general cases, which may be used to obtain the model parameters from experiments.

---

The human brain is arguably the most complex natural object in the known universe. Its complexity, as well as its capabilities, are a product of its roughly 86 billion [1] neurons that make approximately 10^15^ connections [2] with each other. The majority of these connections are chemical synapses, a schematic diagram of which is shown in Fig 1a, with the vesicle-rich pre-synaptic terminal on the left and the post-synaptic terminal with neurotransmitter receptors on the right. In the pre-synaptic terminal, ready-release synaptic vesicles (SVs) packed with neurotransmitters attach to docking sites. The arrival of an AP controls the calcium dynamics which triggers the stochastic fusion of a few SVs (the *quantal content*) with the pre-synaptic membrane, thereby releasing their neurotransmitter cargo in the synaptic cleft as shown. A stochastic train of APs is shown in Fig 1b (top) and the corresponding time dependent quantal content is shown schematically (below). Experimentally the quantal content is obtained as the ratio of the measured peak amplitude of evoked (excitatory or inhibitory) postsynaptic currents [3] to average quantal size. Thus, SV release from pre-synaptic terminals is a fundamental feature of all information processing in the nervous system.

**FIG. 1:**
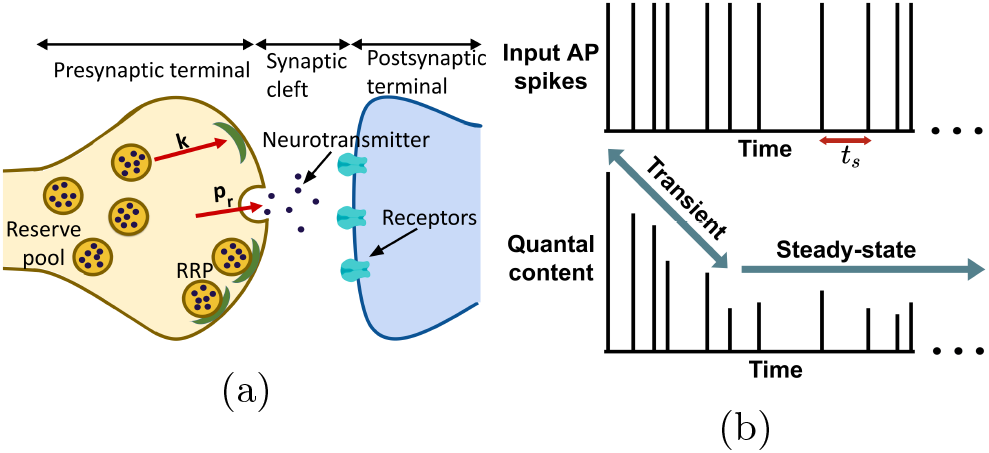
(a) Schematic picture of a chemical synapse. The SVs (circular balls) filled with neurotransmitters (black dots) get docked to sites (in green) and remain ready to release at the presynaptic terminal. Each AP triggers release of some SVs to the synaptic cleft, which reach the receptors in post-synaptic terminal to produce evoked post-synaptic current. (b) A schematic input AP train with random intervals *t*_*s*_ (top), and (bottom) the output quantal content varying with time, shown to reach a steady-state at long times.

Addressing how neurons communicated with another, Castillo and Katz [4] first proposed what they described as a quantal theory of release of certain binomially distributed ‘units’. Over the years micro-structural and electrophysiological studies revealed the relevant biological details, and a picture emerged of spatially distributed pools of neurotransmitter containing SVs in the presynaptic neuron. These are the resting, recycling and ready-release (RRP) pools. SVs from the RRP may release their neurotransmitter cargo on stimulation by an AP through membrane fusion at the pre-synaptic terminal [5–8]. Sustained APs require RRP replenishment, and early theoretical analysis assumed deterministic replenishment, and quantal release [13–15]. Later works developed probabilistic descriptions and stochastic models [16–22]. Empirical observations had shown that the binomial distribution was a good fit to the statistics of quantal release, and this has been the basis of the majority of fluctuation analysis [12, 16, 20, 22–27]. The basic model thus assumes that at the instant of AP stimulation, if there are *n*_−_ docked SVs in RRP and *p*_*rt*_ is the instantaneous release probability, then quantal content *b* (fused and released SVs) is distributed as 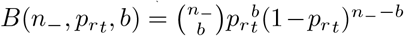 [12, 20–22, 28–30].

As *n*_−_ is itself stochastic, it is generally expected that the actual distribution of quantal content is not a binomial, as would result from summing over the likelihood of random values of *n*_−_. This fact has largely been overlooked in the literature and some works even used binomial with *n*_−_ approximated by its mean value [20– 22, 28, 29, 31, 32]. Systematic formalism treating the docked vesicle number stochastically appeared in a set of works [33–36] – although the focus of the authors was the estimation of the model parameters. While exact relationships between probability of postsynaptic currents after two successive APs were formally written down, the problem could not be solved analytically further because of the presence of multiple nested summations due to the explicit history of the AP train [34].

In this work, we show that in the *steady state* limit after sustained AP stimulation, for uncorrelated stochastic inter-spike intervals (ISI), the full distribution of quantal content *b* as well as its moments becomes *analytically tractable*. The commonly used fixed ISI stimulation follows as a special case of our general result. Although the transient plastic regime (Fig. 1b) is often studied in synapses, the steady state is also of interest [7, 9, 12, 20] and is attained in experimentally reasonable times [37, 38]. Poisson AP stimulation has been experimentally studied [39], and such ISI distributions have been observed in visual cortical neurons [40, 41]. Thus stochastic stimulations are important *in vivo* scenarios. These facts provide motivation for our focus here on steady state quantal content under stochastic stimulations.

In Fig 2, the time dependent number *n*(*t*) of docked SVs in the pre-synaptic neuron is shown schematically. Starting with *n* = *n*_+,*m*−1_ after the (*m* − 1)^*th*^ AP, the number grows stochastically to *n* = *n*_−,*m*_ up to a time *t*_*sm*_, at which the appearance of the *m*^*th*^ AP leads to the fusion and release of *b*_*m*_ vesicles, resulting in a sudden decrease of SVs from *n*_−,*m*_ = *n*_+,*m*_ + *b*_*m*_ to *n*_+,*m*_.

**FIG. 2:**
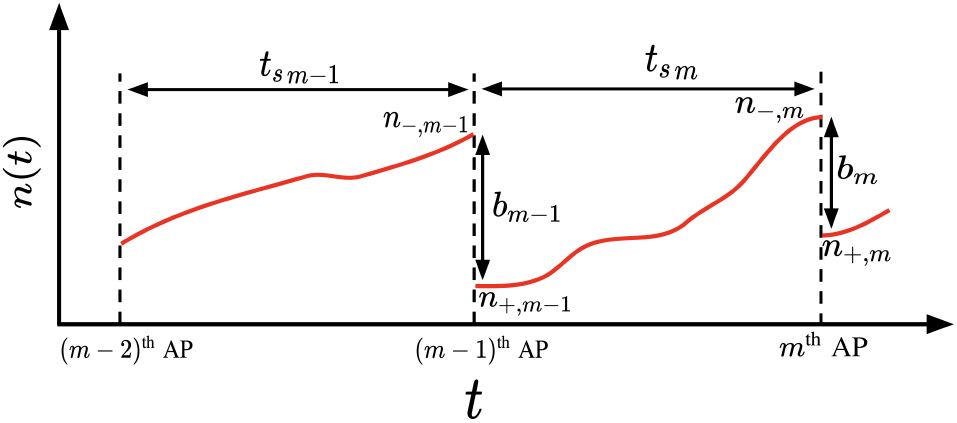
The time variation of docked SV number *n*(*t*) is shown as a function of time over two successive ISIs between the *m* − 2 & *m* − 1 & *m*^*th*^ APs. The respective numbers before an AP (*n*_−_) after an AP (*n*_+_) and quantal content (*b*) are shown.

We assume that docked vesicles may be released probabilistically after an AP but do not otherwise detach. Following the empirical literature [12, 20, 21, 28, 30], as mentioned above the distribution of *b*_*m*_ is 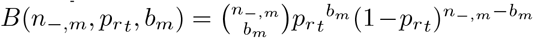 the release probability *p*_*rt*_ is time dependent but saturates at long time. However, *n*_−,*m*_ itself is stochastic and is replenished as follows. We assume *M* is the maximum number of available docking sites, and that docking sites are refilled from the vesicle pool at a rate proportional to the number of empty sites, i.e., *k*_*t*_(*M* − *n*) [20, 22], where *k*_*t*_ is the time dependent docking rate per empty site. The stochastic replenishment of docked SVs from an initial *n*_+,*m*−1_ to some *n* in time *t* is described by the conditional distribution *p*(*n, t*|*n*_+,*m*−1_).

Exact recursive relations may be written between the probabilities 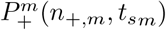and 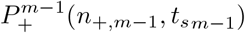 for the SV numbers *n*_+,*m*_ and *n*_+,*m*−1_ after *m*^*th*^ and (*m* − 1)^*th*^ AP respectively – similar to ideas in [33, 34]:

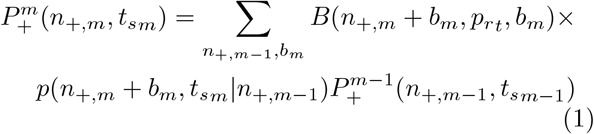

Similarly, the probability of SV number *n*_−,*m*_ just prior to the *m*^*th*^ AP namely 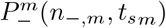 given by:

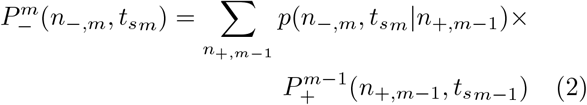

The nested nature of the Eqs. (1) and (2) depending on the full history of successive ISIs make problems of these type analytically very challenging and various computational methods have been developed in this context [33, 34, 36]. In this work we focus on the long time limit after sustained AP stimulation, when interestingly we find that the problem becomes analytically tractable.

Whether the input stimulation is a deterministic AP train [38] or a stochastic one [39], on sustained stimulation (i.e. the index *m* of AP and time is large), a *steady state* [7, 12, 20, 37, 38] is expected to be reached. In this limit, the mean quantal content ⟨*b*⟩ becomes a constant, and its distribution *Q*^*ss*^(*b*) becomes time independent. Analytically obtaining *Q*^*ss*^(*b*) is the main aim of this work. Note that in this *t* → ∞ limit, the time-varying calcium concentration-dependent replenishment rate *k*_*t*_ and release probability *p*_*rt*_ saturates to constant values *k* = *k*_∞_ and *p*_*r*_ = *p*_*r*∞_ (see representative curves in Section 8 of supplementary information (SI) following a model of Ref. [15]). As we solely focus on the steady state, the only two neuronal model parameters we assume in our calculations are *k* and *p*_*r*_. We will show that these can be estimated by comparing experimen-tal data with exact moments of quantal content that we derive.

We study stochastic AP stimulations but assume that the ISIs are uncorrelated, i.e. the joint distribution of two intervals *t*_*sm*_ and *t*_*sn*_ namely *g*_2_(*t*_*sm*_, *t*_*sn*_) = *g*(*t*_*sm*_)*g*(*t*_*sn*_). Under this assumption, for any general stochastic ISI distribution *g*(*t*_*s*_), in steady state, it may be shown (Section 2 of SI) that the Eqs. (1) and (2) become

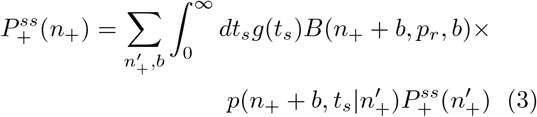

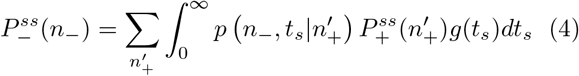

At long times, *m* is large. Therefore, we have used 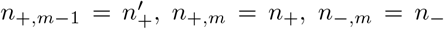, *n*_+,*m*_ = *n*_+_, *n*_−,*m*_ = *n*_−_ and *b*_*m*_ = *b*, as they become history independent random variables. For the case of common experimental interest of fixed ISIs we need to set *g*(*t*_*s*_) = *δ*(*t*_*s*−_*T*). The conditional distribution 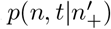 for 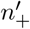to become *n* in time *t* = *t*_*s*_ used in Eqs. (3) and (4), satisfy the Master equation [45]:

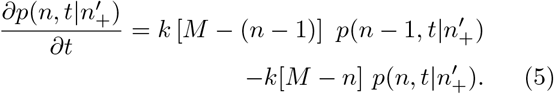

This equation can be solved using the generating function 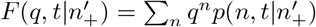 (see SI Section 1 for details), and the solution is:

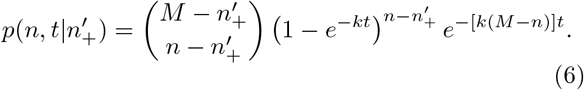

The true steady state distribution of quantal content *Q*^*ss*^(*b*), is obtained by summing its Binomial distribution at every release step over all possible docked random SV numbers *n*_−_:

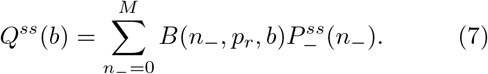

Thus to calculate *Q*^*ss*^(*b*), we need the steady state distribution 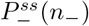, and in turn 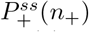because of the relationships in Eqs. (3) and (4). The Eq.(3) has 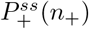 on both sides. Through a lengthy calculation (see details in SI Section 3), we show that the corresponding generation function, namely 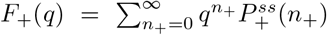 can be obtained using the knowledge of 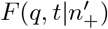, the generating function for 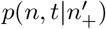 obtained above (Eq. (6)). This generating function leads us to two other relevant gen-erating functions 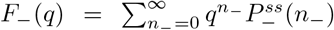 and 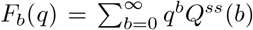 corresponding to the distribu-tions 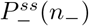and *Q*^*ss*^(*b*). Since the three stochastic variables, *n*_−_, *n*_+_, and _−_*b* are related to each other through the relationship *n*_+_ = *n*_−_ *b*, the different generating functions are also related (see details in SI Section 4):

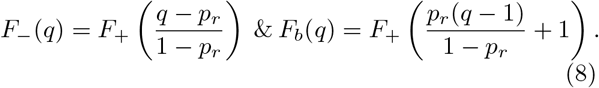

Once we know *F*_*b*_(*q*), we may obtain the desired distribution *Q*^*ss*^(*b*) (see SI Section 4) using the standard relationship 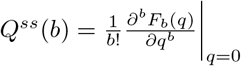 [45]:

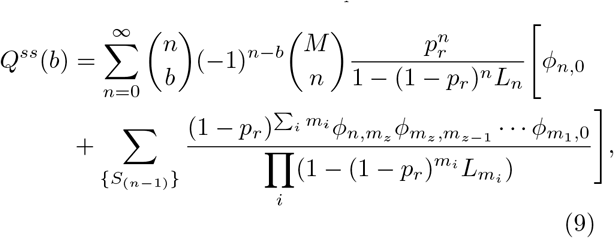

where, 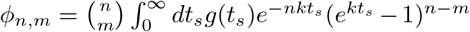 and 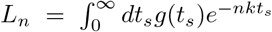. In Eq. (9), {*S*_(*n*−1)_} de-notes the set of all the subsets *S*_(*n*−1)_ = {*m*_*i*_} = (*m*_*z*_, *m*_*z*−1_, …, *m*_1_) with *m*_*i*_ ∈ (1, 2, …, *n* − 1) such that *m*_*z*_ *> m*_*z*−1_ *>* …*> m*_1_. Although the upper limit of the sum is shown up to ∞, due to the combinatorial factor 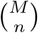, *n* never exceeds *M*. Note that for an arbitrary *g*(*t*_*s*_) there is no reason to expect that *Q*^*ss*^(*b*) in Eq. (9) would reduce to a binomial distribution.

The closed form expressions of the mean and 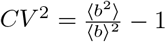 of the distribution in Eq. (9) (see Section 4 in SI) are:

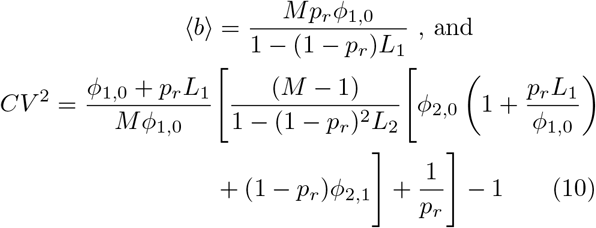

As fixed frequency trains are commonly used in experiments, and Poisson-like statistics have been observed in the ISI distribution from neurons in the visual cortex [40, 41], explicit expressions of *Q*^*ss*^(*b*) in these two special cases are desirable.

For constant ISI *t*_*s*_ = *T*, also called a fixed frequency *f* = 1*/T* AP train, we have *g*(*t*_*s*_) = *δ*(*t*_*s*−_*T*). In SI Section 5(a), we shown that *Q*^*ss*^(*b*) in this case reduces to a binomial distribution:

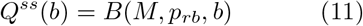

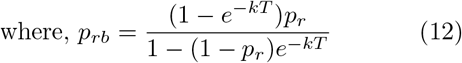

The exact binomial distribution has the parameters *M* and a modified release probability *p*_*rb*_ given by Eq. (12), which depends on the interval *T* and growth rate *k* apart from *p*_*r*_. We do Kinetic Monte Carlo (KMC) simulations to verify our analytical predictions. In Fig 3a, we see that the exact distribution *Q*^*ss*^(*b*) (Eq. (11)) matches perfectly the data from KMC simulations (code in SI) of this process (see the red symbols). Note that Eq. (11) is not equivalent to an approximate binomial often used in the literature *B*(⟨*n*_−_⟩, *p*_*r*_, *b*). In fact we calculate exactly ⟨*n*_−_⟩ = *Mp*_*rb*_*/p*_*r*_ (see SI Section 5(a)), and use this to plot *B*(⟨*n*_−_⟩, *p*_*r*_, *b*) (dashed line) in Fig 3a which deviates significantly. In the large *T* limit, from Eq. (12), *p*_*rb*_→*p*_*r*_ (the release probability in the steady state), as one would expect. The mean and *CV* ^2^ of *b* from Eq. (11) are:

**FIG. 3:**
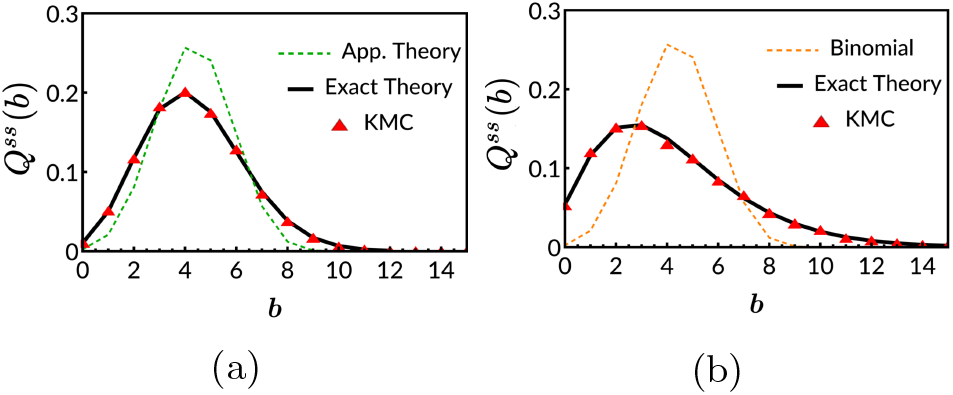
Steady state quantal content distributions for (a) fixed AP arrival times (*f* = 1*/T* = 20 Hz), and (b) exponentially distributed ISIs (rate *f*_0_ = 20 Hz). The common parameters are *M* = 50, *p*_*r*_ = 0.5, *k* = 2 sec^−1^. The exact theoretical curves (thick lines) match the KMC data (red symbols). In (a) an approximate (see text) binomial distribution (dashed line) is shown alongside the exact binomial. In (b) the binomial distribution of Eq. (11) with *T*→1*/f*_0_ (dashed line) deviates from the exact non-binomial distribution.

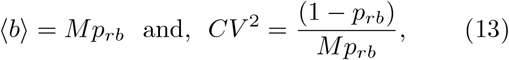

For Poisson AP train, the ISIs are exponentially distributed with 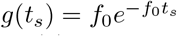. Obtaining *ϕ*_*n,m*_ and *L*_*n*_ in this case, Eq. (9) simplifies (see details in SI Section 5(b)) to yield the following quantal content distribution:

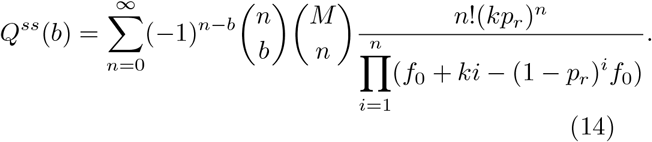

The corresponding mean and *CV* ^2^ for the Poisson case (see Section 5(b) in SI) are:

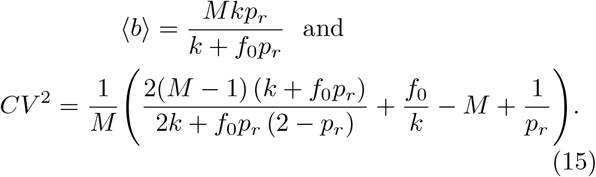

Note that this distribution is no longer a binomial. In Fig 3b we see that the exact formula (solid line) for *Q*^*ss*^(*b*) matches very well with the data obtained from KMC simulation (in symbols). Just to demonstrate that a Binomial approximation would be completely inadequate in this Poisson case, we plot Eq. (11)) with *T* = 1*/f*_0_ where *f*_0_ is the Poisson rate – the deviation is stark.

## Experimental verification and estimation of model pa-rameters

To validate the predictions of our model, we performed electrophysiological experiments (see Section 9 of SI). The comparison between theory and experiment is done in Fig 4. In Figs 4a and 4b, for synapse-1 (left) and synapse-2 (right) respectively, we plot the time-varying fluctuating quantal contents for a single tetanic burst (in blue), as well as the mean quantal content over 20 repeats (in red). In both neurons the mean quantal content reaches a steady-state value beyond 0.8 s (see the yellow box). Considering quantal contents between times 0.8 − 1.0 s over all the 20 histories, we obtained the normalised histogram of the experimental quantal contents – these are shown (in green) in Figs 4c and 4d.

**FIG. 4:**
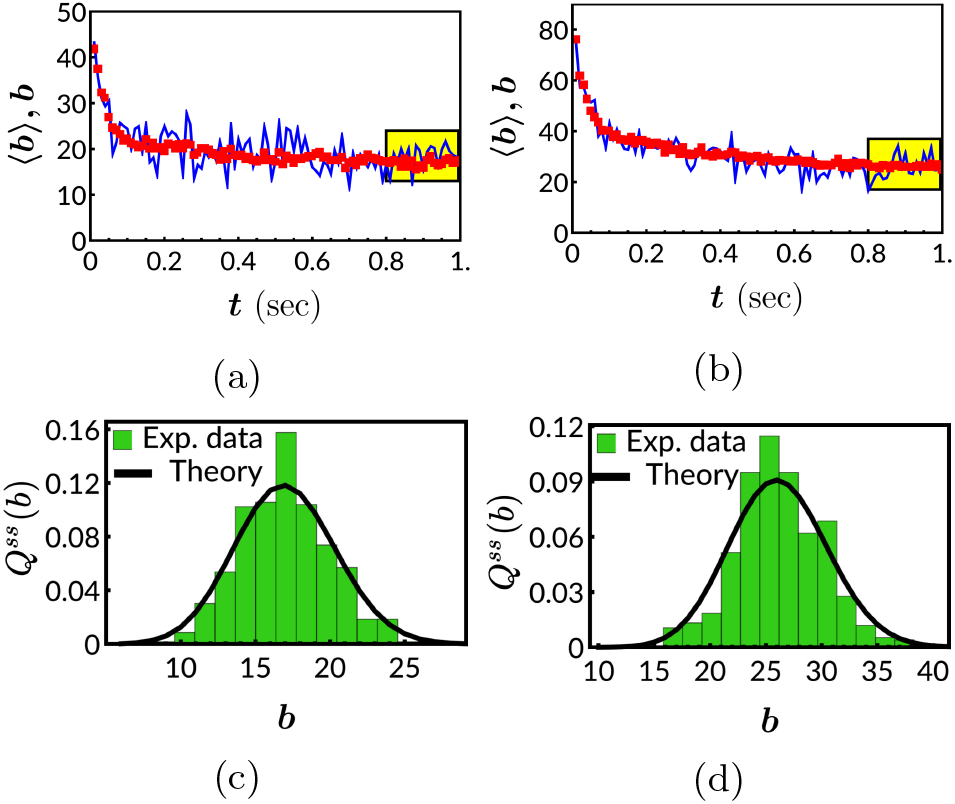
The experimental mean quantal content ⟨*b*⟩ is plotted (in red) against time, for (a) synapse-1 and (b) synapse-2. The blue curves represent one of the 20 experimental realizations of the quantal content. The yellow box shows the part of data we considered to be in the steady state for plotting the experimental histograms of quantal content in (c) and (d) (in green). In (c) and (d), the parameter values used for the exact theoretical distributions (black curves) for synapse-1 are *M* = 50 and *p*_*rb*_ = 0.345 and for the synapse-2 are *M* = 98 and *p*_*rb*_ = 0.266.

After the first AP (at 0 sec), the quantal content *b*_0_ should follow a binomial distribution *B*(*M, p*_*r*0_, *b*_0_), since docking sites should have maximal occupancy.From the 20 experimental histories we obtained the mean and variance of *b*_0_ and using the theoretical formulas of mean = *Mp*_*r*0_ and variance = *Mp*_*r*0_(1 − *p*_*r*0_), we obtained *M* by eliminating *p*_*r*0_. Next we equate the experimental steady-state ⟨*b*⟩ (Figs 4c and 4d) to the theoretical mean ⟨*b*⟩ = *Mp*_*rb*_ (Eq. (13)), and determine the parameter *p*_*rb*_. With these parameters, we plot the exact distribution Eq. (11) (black curves) for the two synapses and they match quite well to the experimental data in Figs 4c and 4d (in green). The *CV* ^2^ automatically turn out quite close: for synapse-1, it is 0.039 (theory) vs 0.032 (experiment), while for synapse-2, the values are 0.028 (theory) vs 0.022 (experiment). Thus we have shown how the full experimental distribution, not just few moments, may be matched to the analytical distribution function. Such comparative study of full experimental and theoretical distributions maybe done for stochastic inputs with Poisson APs (using Eq. 14) or any other *g*(*t*_*s*_) (using Eq. 9) in future.

Note that the moments in the fixed I SI c ase depend on the composite parameter *p*_*rb*_ (Eq. (13)) and not separately on the model parameters *p*_*r*_ and *k*, and hence we cannot estimate them using the fixed ISI data.But that would not be the case for Poisson APs – see Eq. (15)) where the explicit dependence on *p*_*r*_ and *k* appear. Thus future experimental data on Possion APs may be used to estimate the model parameters. In fact this may be done for stochastic inputs with any other *g*(*t*_*s*_)– Eq. (10) may be used in such cases.

Exact results are rare and desirable in biological contexts. Here, we analyze the basic model of synaptic transmission between neurons applicable to all chemical synapses. Although we look at the restricted regime of steady state and uncorrelated stochastic AP inputs, we provide full exact general distributions including moments. We also show how the results may be used to compare with experimental data and indicate how model parameters may be estimated in future.

## Acknowledgements

DD acknowledges IIT Bombay for funding. KR thanks IIT Bombay for Institute Ph.D. fellowship. AS acknowledges support by NIH/NIDCD grant R01DC019268. EF acknowledges support by Bundesministerium für Bildung und Forschung grant 01GQ2001.

## 1 Probability distribution of docked vesicle number *n*between two consecutive action potentials

Let 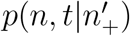 denote the probability distribution of the number *n* of docked (ready to release) vesicles at time *t*, starting from 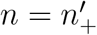 at *t* = 0 (i.e. from the instant of the last action potential (AP)). Thus *t* denotes any time up to the next AP, i.e. *t < t*_*s*_ where *t*_*s*_ is a time when the next AP arrives. In the interval *t* to *t* + *dt*, the number of docked vesicles may increase from *n* − 1 to *n* with the probability *k*(*M* − (*n* − 1))*dt*. Here *k* is the growth rate per empty docking sites, and *M* is maximum number of docking sites. Consequently, the master equation for the time evolution of 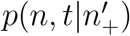 between two successive APs is:

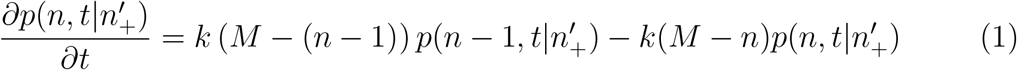

We define a generating function for the above probability distribution:

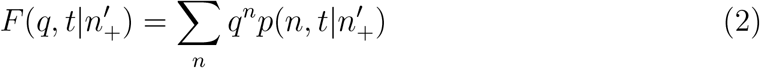

Here the limits of the above sum are 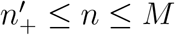. Otherwise, we assume 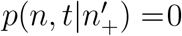 if 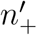falls outside this range. Multiplying Eq. (1) by Σ_*n*_ *q*^*n*^ on both sides, we get a PDE for the generating function *F* :

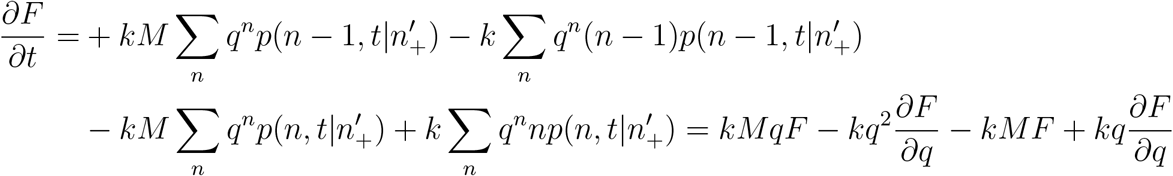

Rearranging the terms, we have,

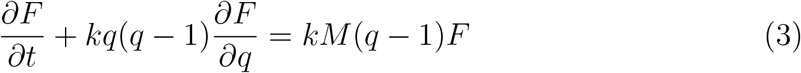

We use the method of Lagrange characteristics to solve the Eq. (3). The result is:

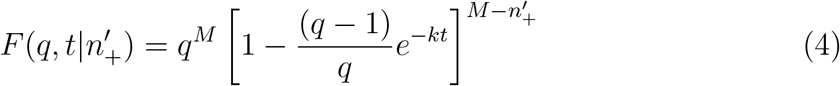

We do binomial expansion of Eq. (4) to get

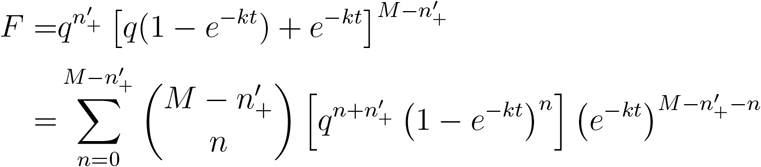

and relabelling 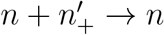 we get

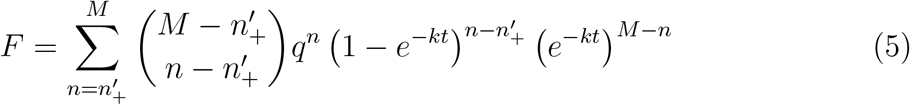

Next we can compare Eq. (5) with Eq. (2) to get the expression of 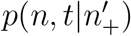:

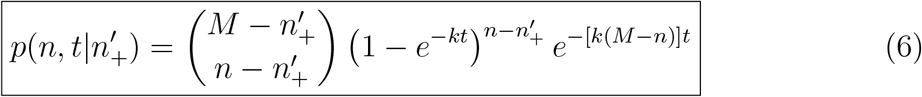

for 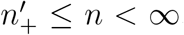. Note that for *n > M*, the probability goes to zero automatically in the above expression.

## 2 Self-consistent equation for the quantal content distribution 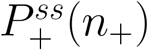 in the steady state

Let us denote the time intervals between (*m* − 1)^*th*^ AP and *m*^*th*^ AP by *t*_*sm*_ and between the (*m* − 2)^*th*^ AP and (*m* − 1)^*th*^ AP by *t*_*sm−*1_. We first relate the probability 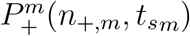 of having *n*_+,*m*_ number of docked vesicles after the *m*^*th*^ AP, to the probability 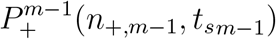 of having *n*_+,*m−*1_ number of vesicles just after the (*m*−1)^*th*^ AP. To do so, we need to sum over all possible paths beginning at *n* = *n*_+,*m−*1_ after the (*m* − 1)^*th*^ action potential and ending at *n* = *n*_+,*m*_ after the *m*^*th*^ AP.

The sum involves the product of three probabilities due to a sequence of steps: (i) initial 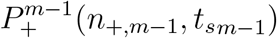, then (ii) the probability *p*(*n*_+,*m*_ + *b, t*_*sm*_|*n*_+,*m−*1_) that *n*_+,*m−*1_ rises to *n*_+,*m*_ + *b* in time *t*_*sm*_, and (iii) finally the probability that at *t*_*sm*_, the number *n*_+,*m*_ + *b* of docked vesicles suddenly reduce to *n*_+,*m*_ due to the AP triggered vesicle release, with the associated quantal content *b* being binomially distribution *B*(*n*_+,*m*_ +*b, p*_*r*_, *b*). After summing over all possible initial number *n*_+,*m−*1_ and quantal content *b* we have,

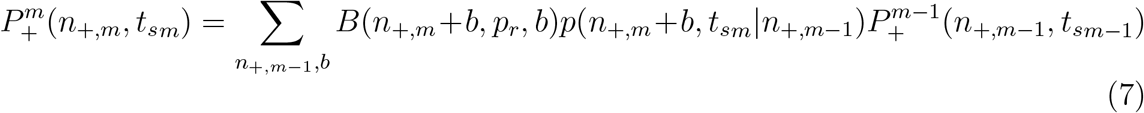

We may follow the same procedure for 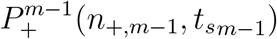 and so on, and build up a hierarchy of relations for any interval between two APs. The problem remains hard to solve as result for every interval depends on the history of the previous interval. We are particularly interested in the steady state attained after several APs appear, i.e. *m* ≫ 1 and the output quantal contents statistically stabilise. We integrate Eq. (7) over the joint probability distribution *g*_2_(*t*_*sm*_, *t*_*sm−*1_) of *t*_*sm*_ and *t*_*sm−*1_. We get

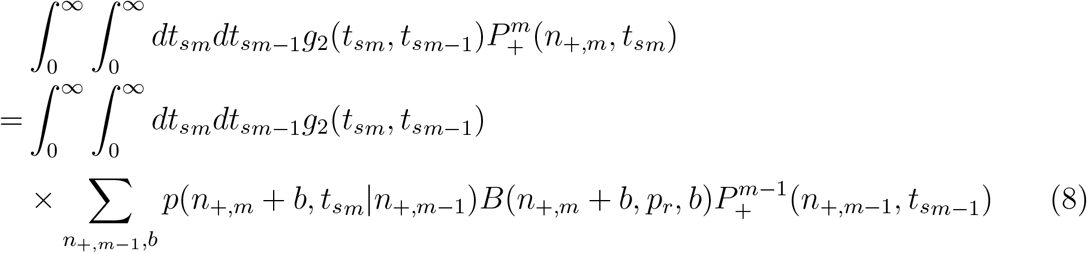

If the interspike intervals (ISIs) are not correlated, we would have *g*_2_(*t*_*sm*_, *t*_*sm−*1_) = *g*(*t*_*sm*_)*g*(*t*_*sm−*1_), where *g*(*t*_*sm*_) and *g*(*t*_*sm−*1_) are the normalised distributions of *t*_*sm*_ and *t*_*sm−*1_ respectively. Then,

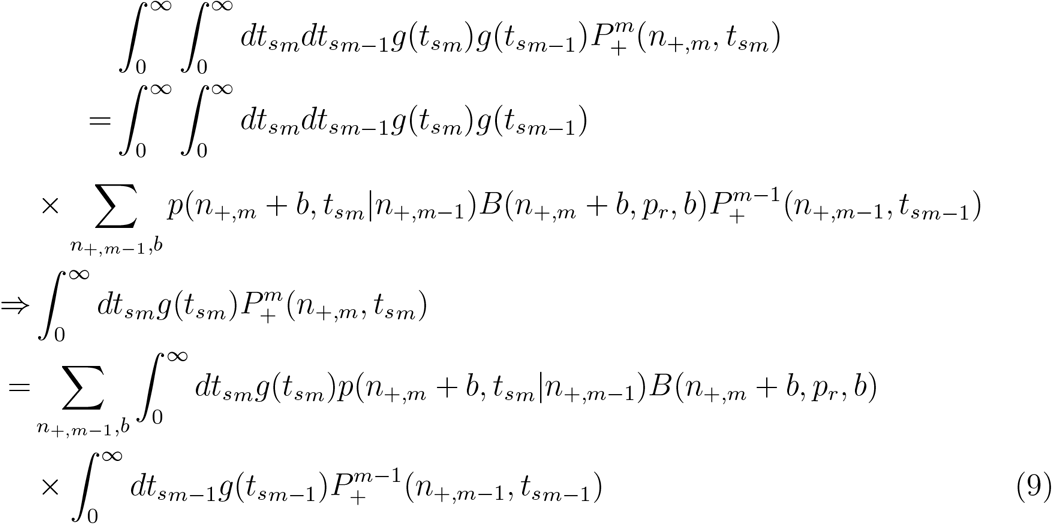

In the long time limit, the steady state probability distribution of vesicle number after any AP is 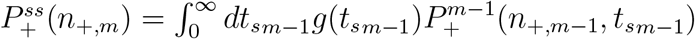, which does not depend any more on the exact history of the spike train. Dropping the subscripts *m* (in the steady state), and setting *t*_*sm*_ = *t*_*s*_ and 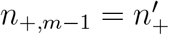, the Eq. (9) becomes

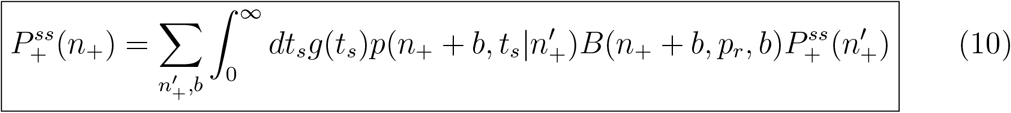

Note that this is a self-consistent equation for the unknown 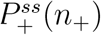, when we know *g*(*t*_*s*_), *B*(*n*_+_ + *b, p*_*r*_, *b*) and 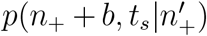 (Eq. (6)). This equation appears in the main manuscript. We analytically treat it in the next section.

## 3 Generating function of the distribution 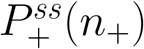 in the steady state

We define a generating function of 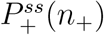as follows and solve it in this section:

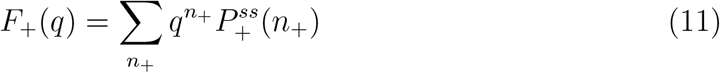

In Eq. (10), the limits of the summation are 0 ≤ *b* ≤ *M* − *n*_+_ and 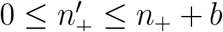. Since anyways, 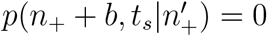 outside the limits (see Eq. (6)), we may as well use the range 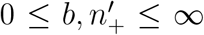. Multiplying Eq. (10) by 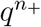 and summing over *n*_+_ (with 0 ≤ *n*_+_ ≤ ∞), we have:

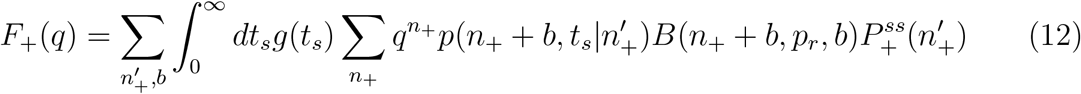

We introduce a new variable *ñ*_+_ = *n*_+_ +*b* with *b* ≤ *ñ*_+_ ≤ ∞. Note that *B*(*ñ*_+_, *p*_*r*_, *b*) = 0 for *ñ*_+_ *< b*, so we may keep the range 0 ≤ *ñ*_+_ ≤ ∞.

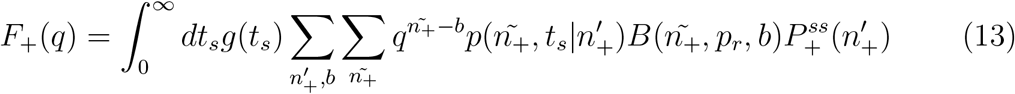

Summing over *b* after substituting 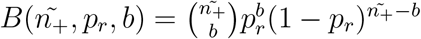, we have

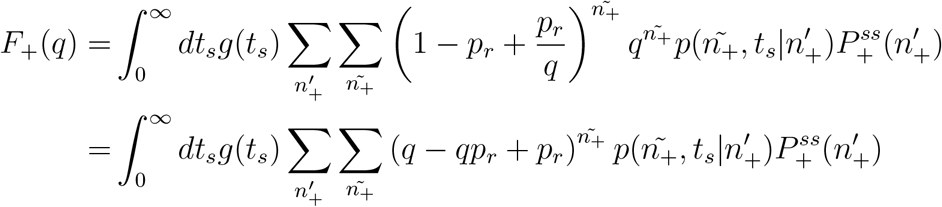

Doing summation over *ñ*_+_ and using the definition of the generating function *F* (from Eq. (2)), we get

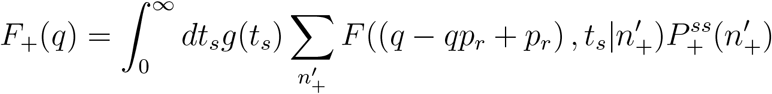

Defining *h*(*q*) = (*q* − *qp*_*r*_ + *p*_*r*_) and using Eq. (4) for the explicit form of function *F* :

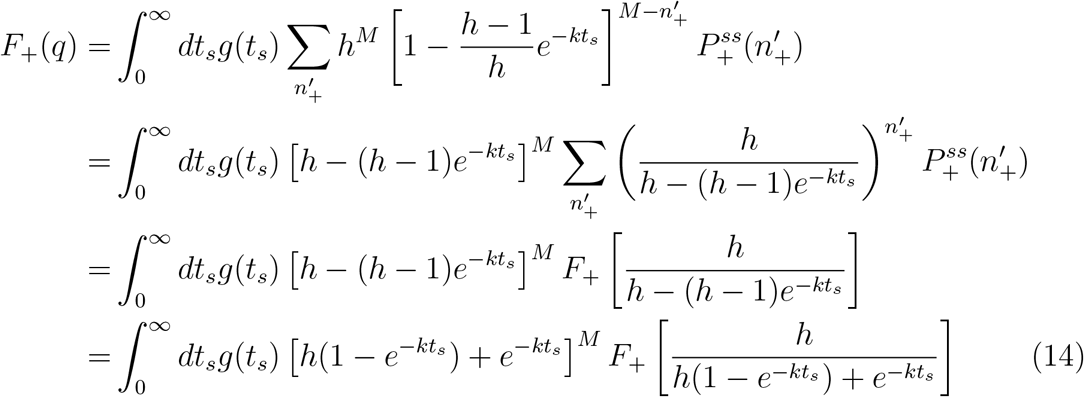

In the above we have used the definition of *F*_+_ from Eq (11). In Eq. (14) we have a self-consistent integral equation for the function *F*_+_ with two different arguments, and involving an integration over the inter-spike interval *t*_*s*_. Clearly, it is difficult to solve the equation directly for *F*_+_(*q*). But as we show below, *F*_+_(*q*) may be obtained as a series expansion around *q* = 1, that is

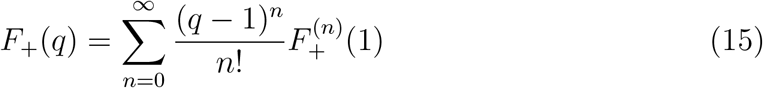

where the series coefficients 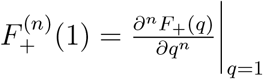. The procedure of obtaining these coefficients are as follows.

Taking derivative with respect to *q* on both side of Eq. (14), we have

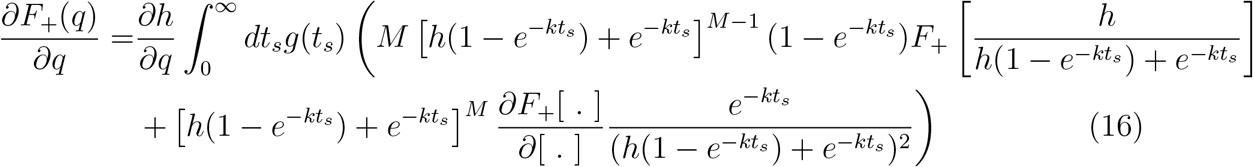

where [·] is a short hand for 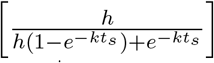. Since 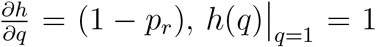, [.]|_q=1_ = 1, F+(1) = 1,and 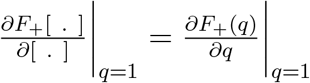,we have

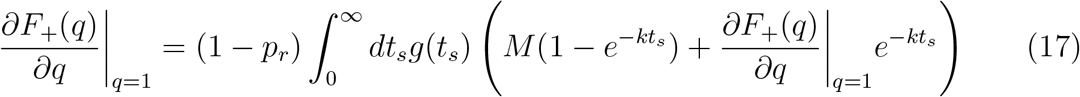

Defining the quantities *Ψ*_*n,m*_ and *L*_*n*_ as follows,

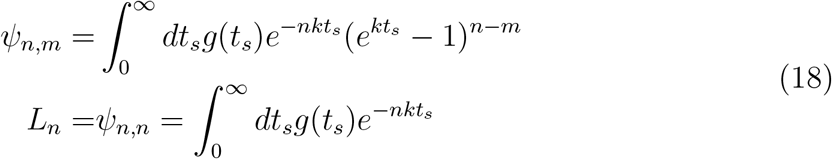

we have

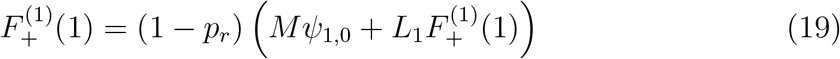

where we have set 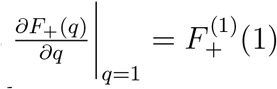 according to the general definition of the *n*-th series coefficient above.

Next we may take derivative of Eq. (16) with respect to *q* and set *q* = 1 and obtain an equation for 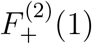. We find that it is linearly coupled to 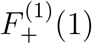 as follows:

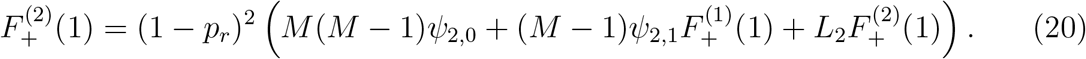

We may follow the same procedure to obtain equations for the higher derivatives. It is clumsy to do it manually, so we use Mathematica for this. We use symbolic derivatives of Eq. (16) and automatic replacement of terms by the *Ψ* functions using the *Replace* syntax. The result are the equations for the next two series coefficients.

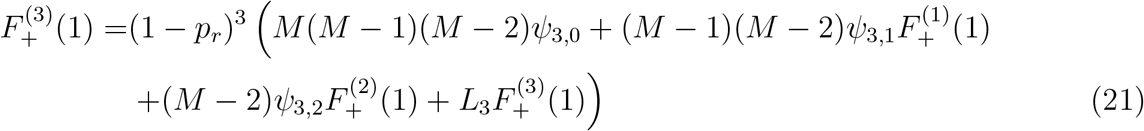

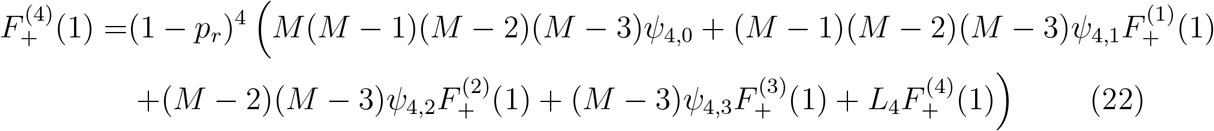

Eqs. (19) -(22) relate the four derivates 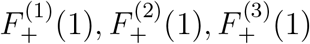 and 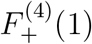 linearly. So, it is easy to obtain solutions by inverting a 4 × 4 matrix, using Mathematica. The explicit expressions of the four coefficients are

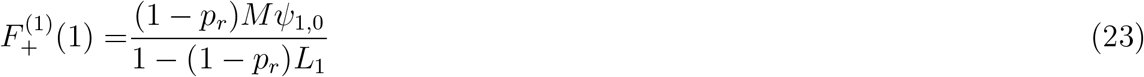

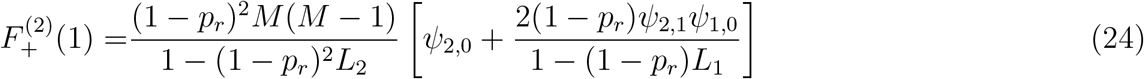

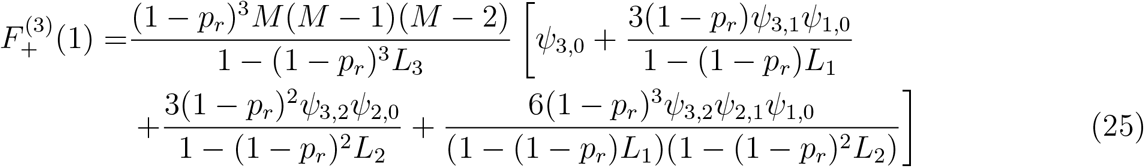

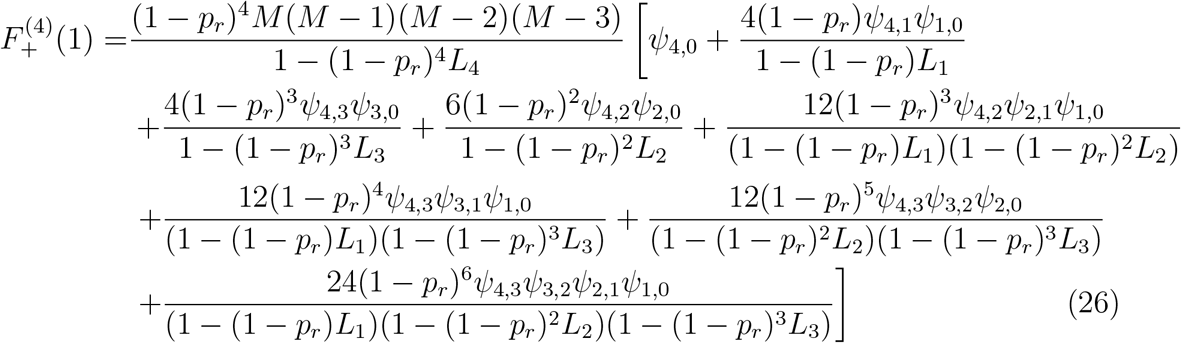

In the above expressions, Eqs. (23) -(26), we notice the following patterns. The terms outside the square brackets are of the form 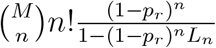 for the *n*-th coefficient. Except for the first term, the denominators of the terms inside the square brackets for the *n*-th coefficient involve factors (1 − (1 − *p*_*r*_)^*j*^*L*_*j*_) with *j*≠*n* or 0. The most crucial fact is that each such term may be associated with a set of integers which are subsets of the set of integers {1, 2, …, *n* − 1}. For example, a subset of *z* ≤ (*n* − 1) positive integer elements is *S*_(*n−*1)_ = (*m*_*z*_, *m*_*z−*1_, …, *m*_1_) = {*m*_*i*_} with *i* = *z, z* − 1, …, 1, and every *m*_*i*_ ∈ {1, 2, …, *n* − 1} and *m*_*z*_ *> m*_*z−*1_ *>* …*m*_1_. The power of (1 − *p*_*r*_) in the numerator of each term is equal to 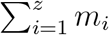. The subscript of *Ψ* factors are formed by pairs of successive integers drawn from the ordered array of numbers: *n, m*_*z*_, *m*_*z−*1_, …, *m*_1_, 0. The numeric factors in any numerator can be absorbed into the *Ψ* factors by introducing the quantities 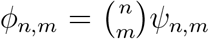. Putting together all these observations of the patterns, we may write down a general form of 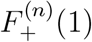 as follows:

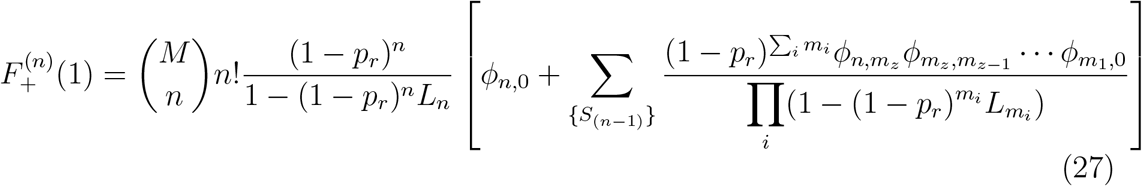

where the sum over {*S*_(*n−*1)_} runs over all such subsets as *S*_(*n−*1)_ mentioned above.

Using Eq. (27) in Eq. (15), we can finally write down the expression of *F*_+_(*q*) as a series expansion around *q* = 1,

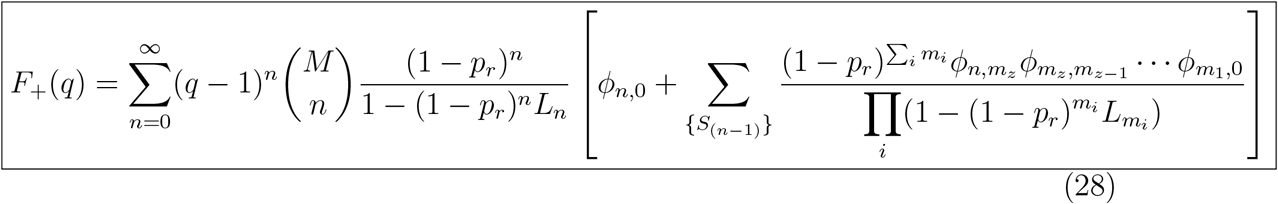

## 4 Relations between the generating functions, and the quantal content distribution *Q*^*ss*^(*b*)

Let us denote the probability distribution of having *n*_*−*_ number of docked vesicles before an AP in the steady state by 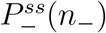. It is easy to see from similar arguments as in Section 2, that 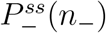will be related to the three distributions: 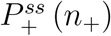 (to have an initial number), *p* (*n*_*−*_|*t*_*s*_, *n*_+_) (for docked vesicles to build up), and *g*(*t*_*s*_) (for AP to appear). Thus

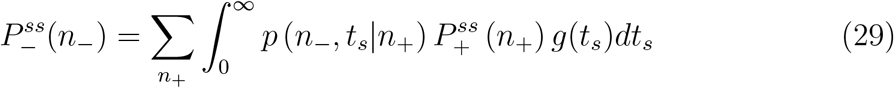

We define its generating function as

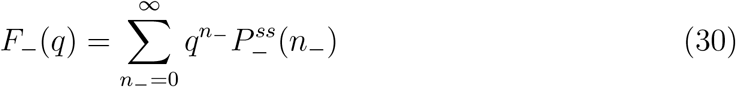

The ultimate object of our interest, namely the distribution of quantal content *b* in the steady state, is related to the distribution 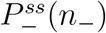 as:

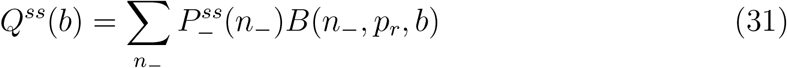

Note that the the crucial fact is that we are averaging the well-known binomial distribution now, over the stochastically generated docked vesicle number *n*_*−*_ just prior to an action potential. As we would see, the resultant averaged distribution would be different from the original binomial. The generating function corresponding to *Q*^*ss*^(*b*) is defined as

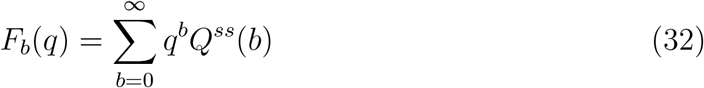

We have four distributions *p* (*n*_*−*_, *t*_*s*_|*n*_+_), 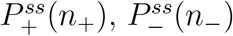, and *Q*^*ss*^(*b*) and four corresponding generating functions *F, F*_+_, *F*_*−*_ and *F*_*b*_, respectively. Of these, we have already calculated *F* (Eq. (4)) and *F*_+_ (Eq. (28)). We would now proceed to relate the generating functions *F*_*−*_ and *F*_*b*_ to *F*_+_. Once *F*_*b*_ is known explicitly using the expression of *F*_+_, we would then obtain *Q*^*ss*^(*b*) by taking suitable derivatives of *F*_*b*_(*q*) with respect to *q*.

Using Eq. (29) in Eq. (30) we get,

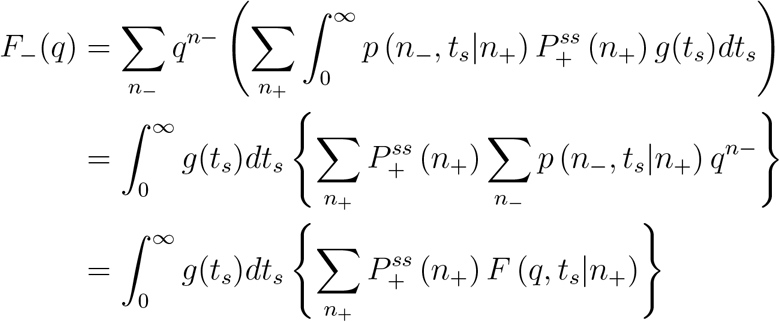

In the above we have used the definition of the function *F*. Substituting the explicit form of *F* (*q, t*_*s*_|*n*_+_) from Eq. (4), we have

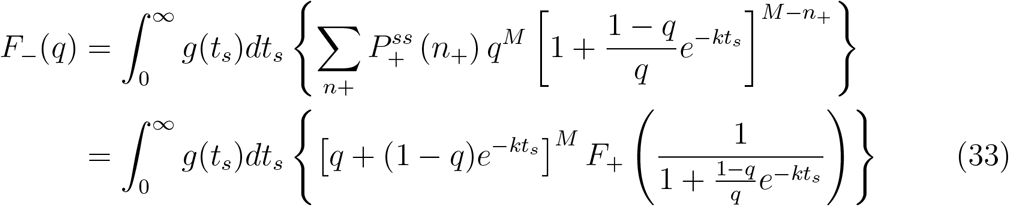

In the above we have used the definition of *F*_+_.

Substituting 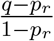 in place of *q* in Eq. (14), and noticing 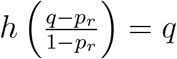, we get

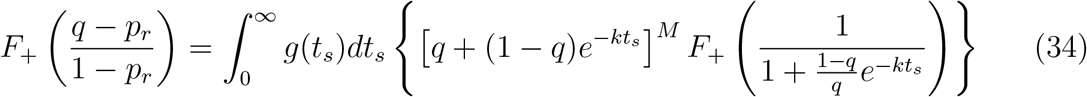

Comparing Eqs. (33) and (34), we immediately get,

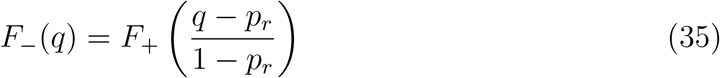

Substituting Eq. (31) in Eq. (32) with the binomial distribution *B*(*n*_*−*_, *p*_*r*_, *b*) written explicitly, we have

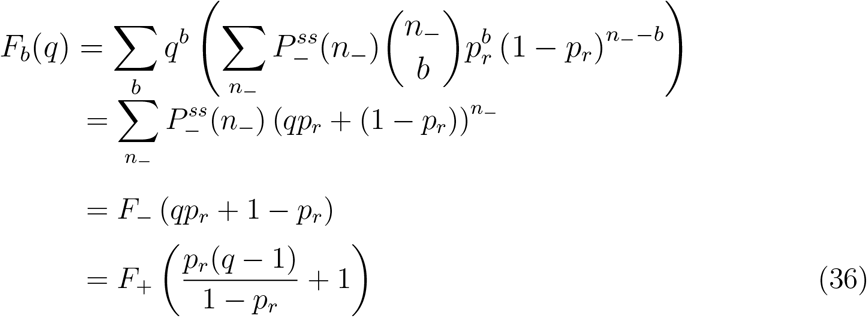

Here we have used the definition of the function *F*_*−*_, and the relation of *F*_+_ to *F*_*−*_ from Eq. (35). Using the explicit form of *F*_+_ from Eq. (28), the Eq. (36) gives,

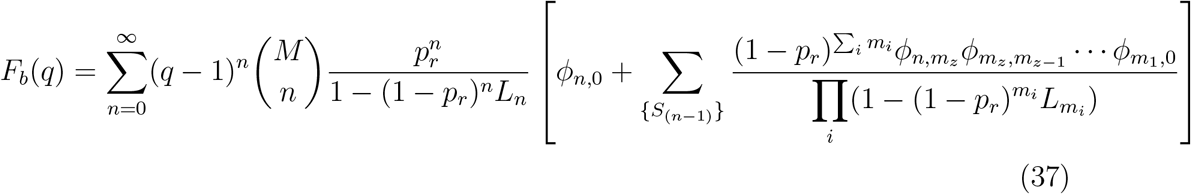

We may invert Eq. (32), to obtain the quantal content distribution *Q*^*ss*^(*b*) from successive derivatives of *F*_*b*_(*q*) as follows:

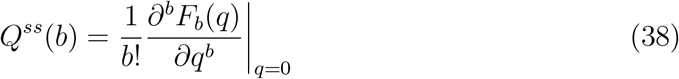

Note that *q* is prese1nt in *F*_*b*_(*q*) only through the factor (*q* − 1)^*n*^ in Eq. (37). Using the identity 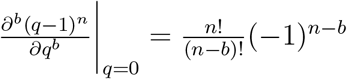, we obtain our final desired result

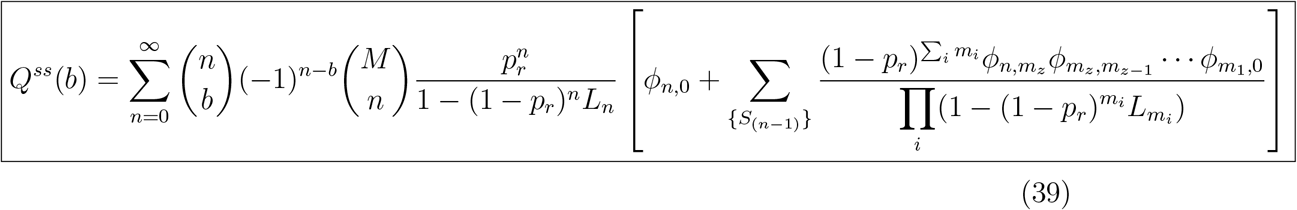

### Moments

(i)For the general distribution of ISIs, following Eq. (39), the mean quantal content ⟨*b*⟩ is given by

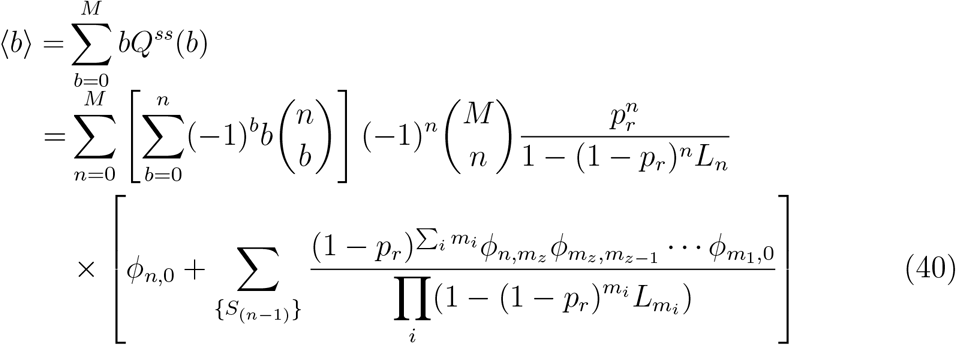

Note that

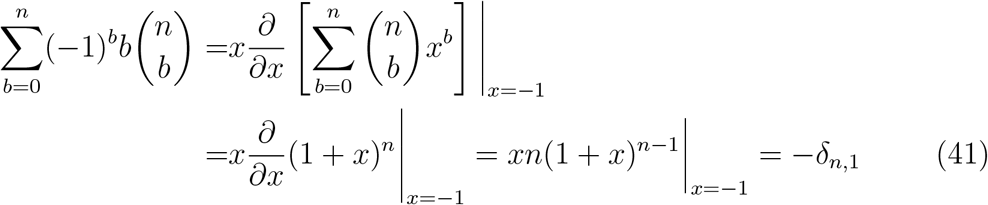

Substituting Eq. (41) in Eq. (40), we have

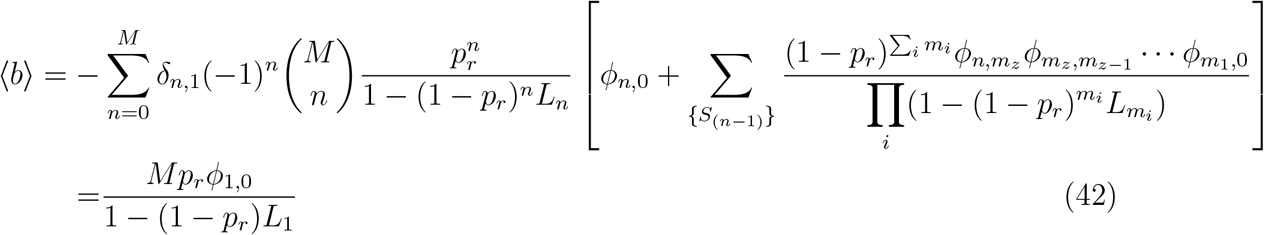

(ii)Similarly, the second moment ⟨*b*^2^⟩ is given as:

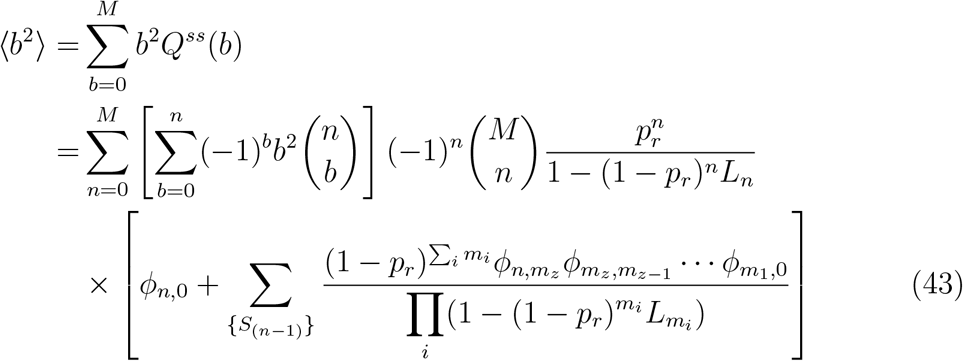

Where,

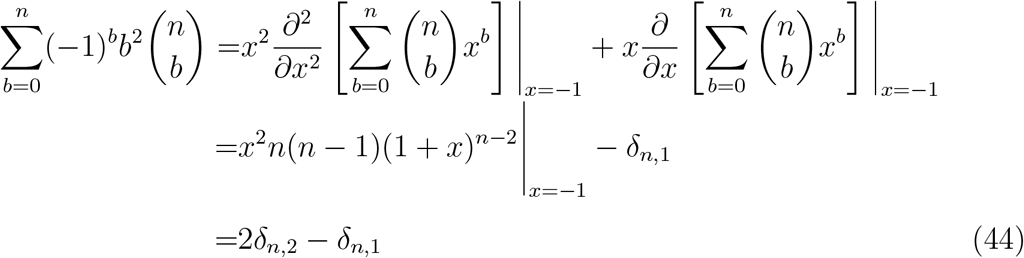

Substituting Eq. (44) in Eq. (43), we have

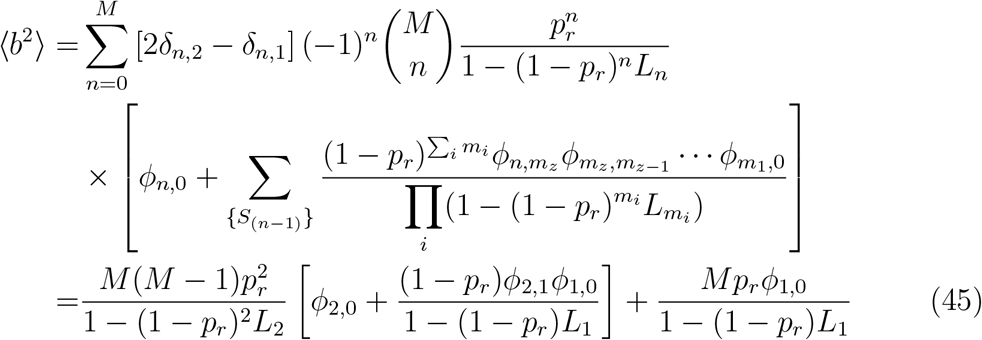

Using Eqs. (42) and (45), *CV* ^2^ can be easily obtained as

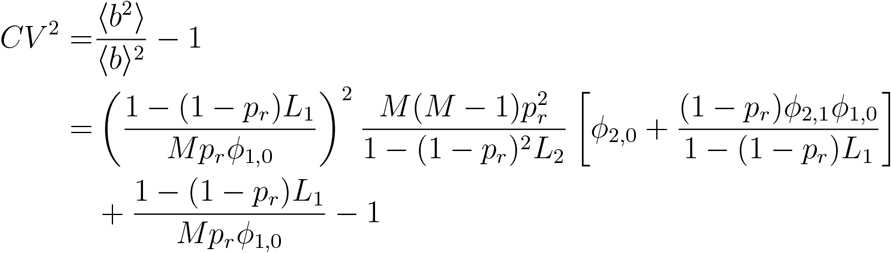

Note that 1 − *L*_1_ = *ϕ*_1,0_. So, we have

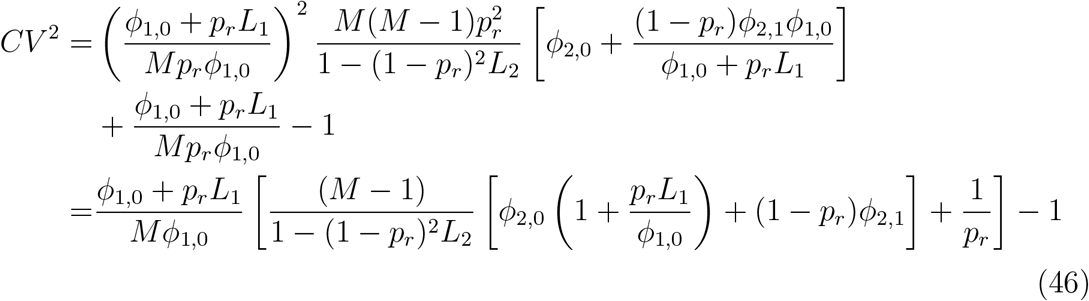

## 5 The distribution *Q*^*ss*^(*b*) for two special cases of inter AP interval distribution *g*(*t*_*s*_)

### (a)The deterministic case – APs at a fixed frequency *f* = 1*/T*

Assuming the APs arrive after fixed time duration *T*, the corresponding distribution *g*(*t*_*s*_) = *δ*(*t*_*s*_ − *T*). This is most often the pattern of input spike train used by experimentalists to excite neuronal response at a fixed frequency *f* = 1*/T*. We may calculate using Eq. (18),

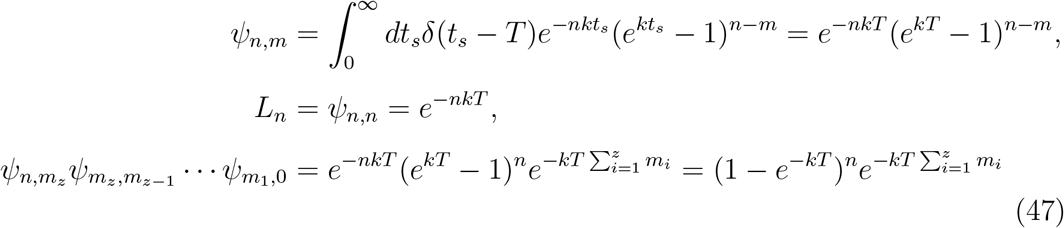

Substituting these expressions in the Eq. (39), and recalling 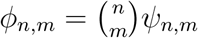, we have

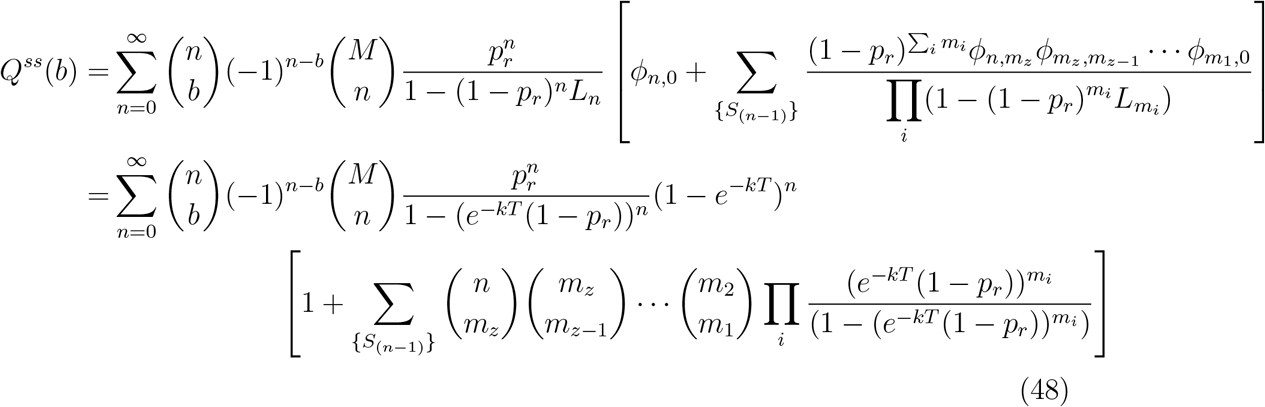

It is important to remind that {*S*_(*n−*1)_} is the set of all the subsets of the set {1, 2, …, *n* − 1}. To proceed further, we need to use the following identity (for |*x*| *<* 1) – for proof see Section 6:

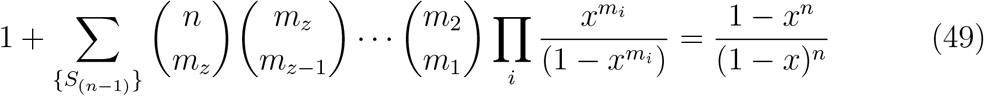

Using the identity (Eq. (49)) in Eq. (48), the expression simplifies to give,

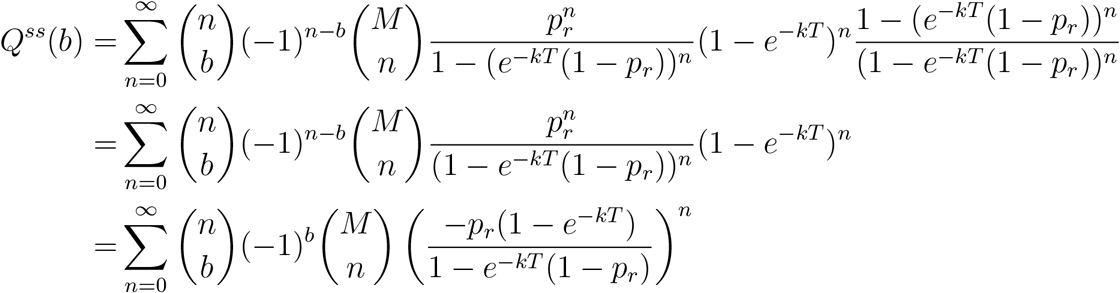

Further using the identity, 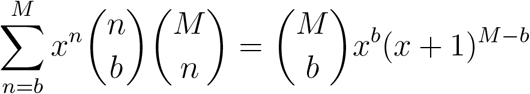 we finally get

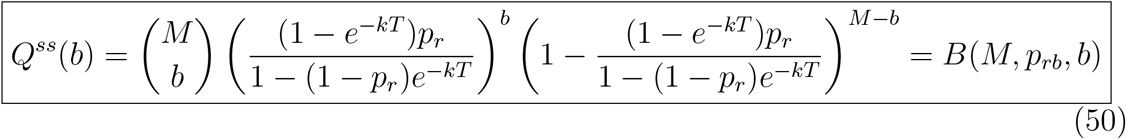

Hence, the steady state quantal content distribution for the deterministic *g*(*t*_*s*_) is still a binomial distribution, but with the parameters *M* (maximal docked vesicle number) and an effective release probability *p*_*rb*_ (different from *p*_*r*_) given by:

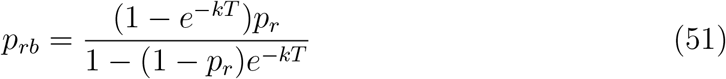

In order to compare the exact distribution (Eq. (50)) with the approximate binomial distribution *B*(⟨*n*_*−*_⟩, *p*_*r*_, *b*) where ⟨*n*_*−*_⟩ is the mean number of docked vesicles just before the AP spikes, we need to calculate an expression for ⟨*n*_*−*_⟩. It is easy to calculate ⟨*n*_*−*_⟩ from the generating function *F*_*−*_(*q*) as 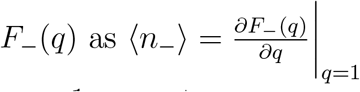.

Taking derivative of Eq. (35) and setting *q* = 1, we get

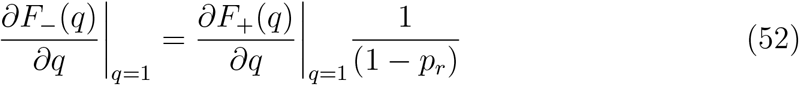

If we take the derivative of Eq. (28) with respect to *q* at *q* = 1, only *n* = 1 term in the summation survives. Eq. (52) will become

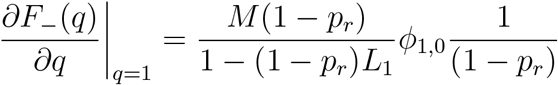

Substituting the forms of *L*_1_ and *ϕ*_1,0_ from Eq. (47), we get

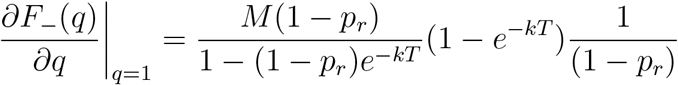

Hence,

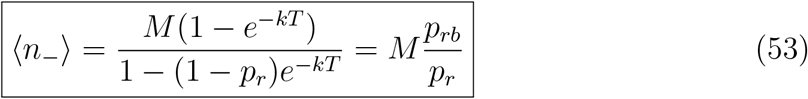

In the above we have used the expression of *p*_*rb*_ from Eq. (51). The Eq. (53) implies that the mean ⟨*b*⟩ of the exact binomial and the approximate binomial are equal, i.e. *Mp*_*rb*_ = ⟨*n*_*−*_⟩*p*_*r*_. But the variance of the exact distribution is *Mp*_*rb*_(1 − *p*_*rb*_), different from that of the approximate binomial ⟨*n*_*−*_⟩*p*_*r*_(1 − *p*_*r*_).

### (b) Poisson distributed AP arrival times

APs may arrive at intervals which may be noisy. A simple way to incorporate stochasticity in the input spike train is to assume that they arrive at time intervals which are exponentially distributed with 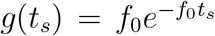. Note that the mean as well as characteristic spike interval is 1*/f*_0_. In this case using Eq. (18),

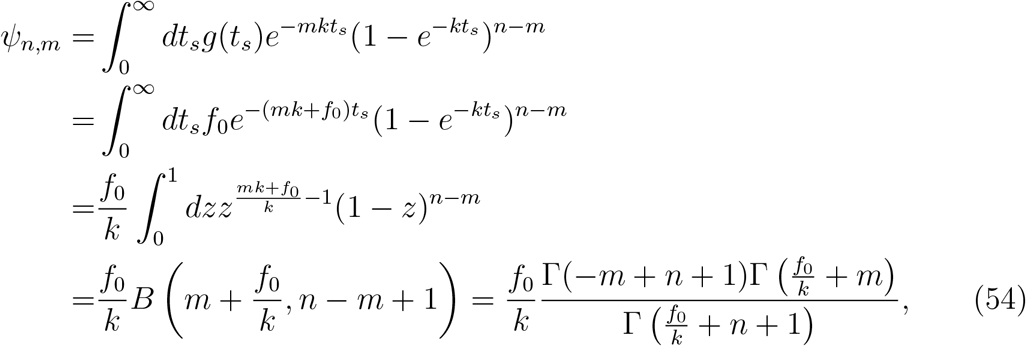

where we have used Beta function *B*(*a, b*) = Γ(*a*)Γ(*b*)*/*Γ(*a* + *b*), and then

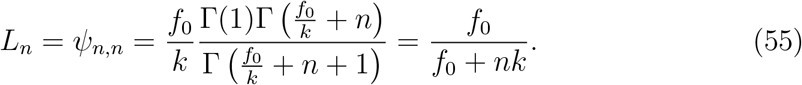

Moreover using Eq. (54), we have

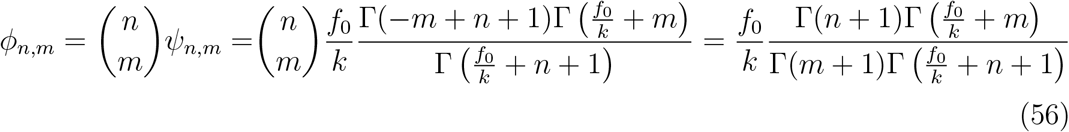

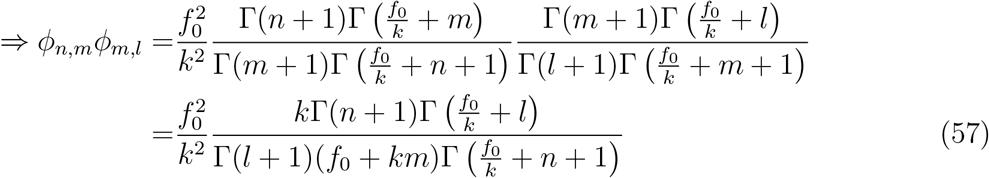

The above Eq. (57) may be extended to get the following product:

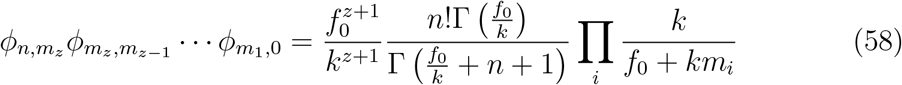

Substituting Eq. (58) into Eq. (39), we have

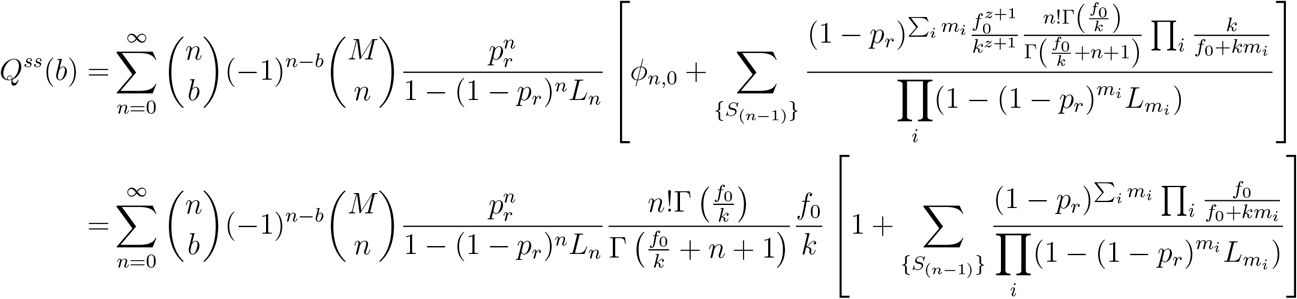

From Eq. (55), substituting 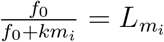, we have

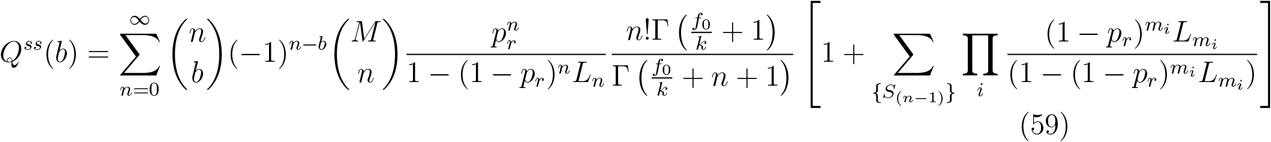

We may simplify the expression above by using the following identity valid for any arbitrary function *f* – for proof see Section 7:

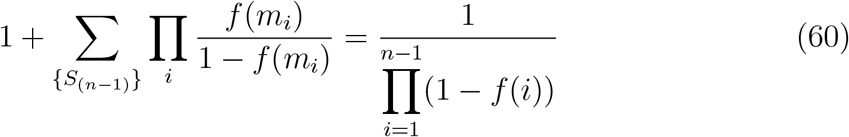

Using Eq. (60) in Eq. (59), we get

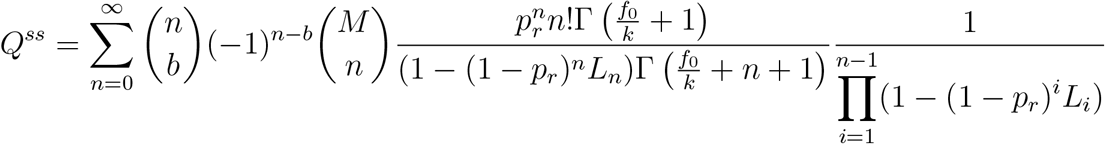

Substituting back 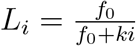,we have

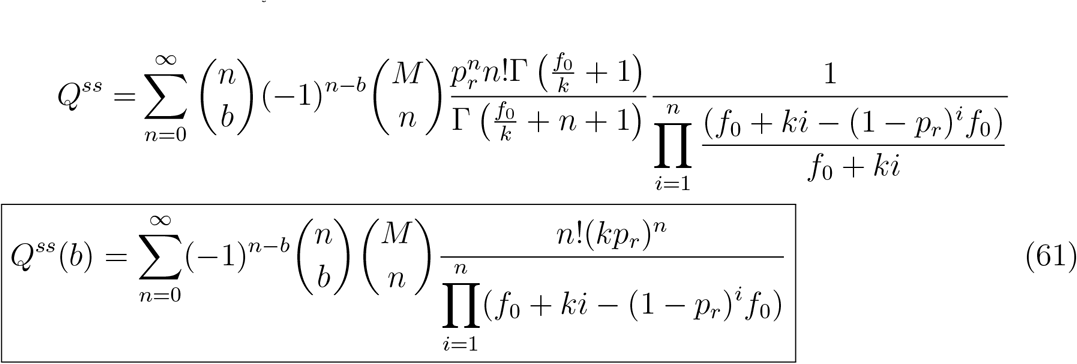

Thus the steady state quantal content distribution for the spike intervals drawn from a Poisson processes is not a Binomial, but instead the above distribution in Eq.

### Moments

(i)For the exponentially distributed ISIs, the mean quantal content ⟨*b*⟩ can be obtained by substituting 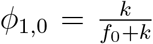 and 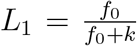 from Eqs. (56) and (55) respectively, into the general ⟨*b*⟩ expression (Eq. (42)) as follows

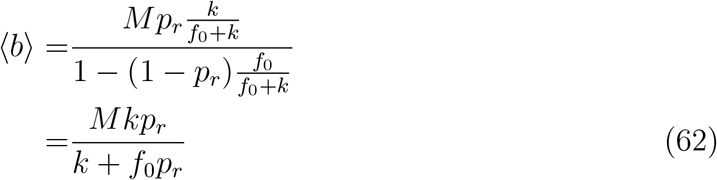

(ii)Similarly, the *CV* ^2^ can be obtained by substituting the expressions: 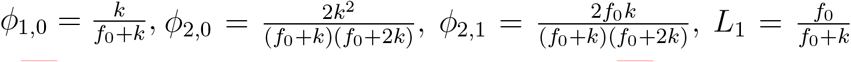 and 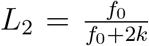from Eqs. (56) and (55), into the general *CV* ^2^ expression (Eq. (46)) as follows

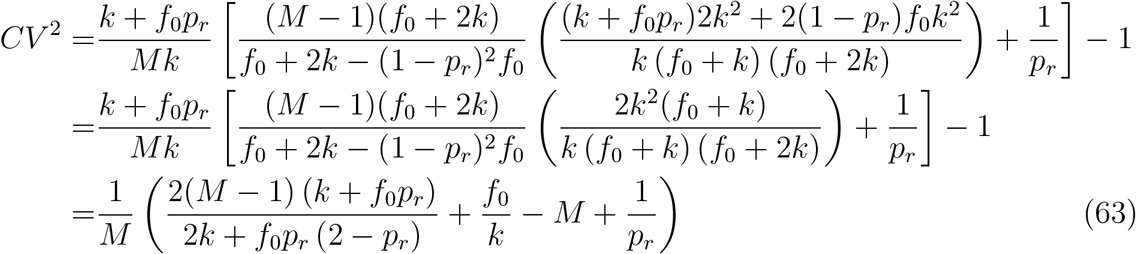

## 6 Proof of the identity in Eq. (49)

The identity can be proved using the principle of strong mathematical induction as follows.

We first verify the identity for *n* = 1 and 2 explicitly. For *n* = 1, *LHS* of Eq. (49) will be simply equal to 1 as the summation term does not contribute anything. The *RHS* is also equal to 1, so the identity is true for *n* = 1. For *n* = 2, {*S*_(1)_} = {{1}}. So, the *LHS* of Eq. (49) will be equal to 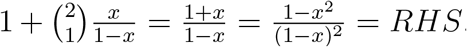. Hence, we have verified the identity for *n* = 2 as well.

Let us assume that the identity is true for all integer values of *n* ∈ [1, *k*]. We would then show below that it is true for *n* = *k* + 1. For *n* = *k* + 1 we have:

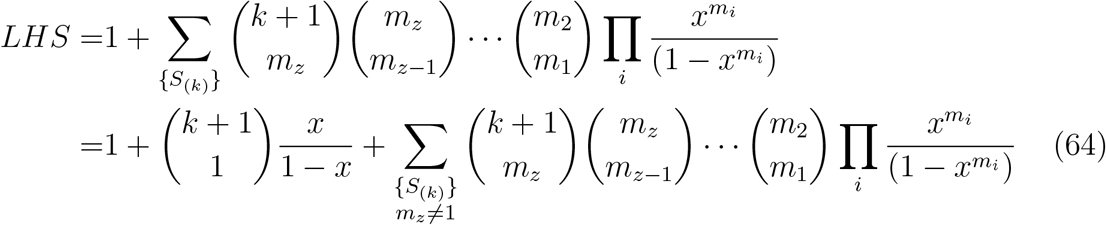

where in the last step, we have separated out the contribution of *m*_*z*_ = 1 in the summation. The crucial step in our proof comes from the idea that the subsets in the set {*S*_(*k*)_} (with *m*_*z*_ ≠1) can be clubbed into different groups, such that the largest element in all the subsets in a particular group are equal. There will be *k* − 1 such groups as the maximum element *m*_*z*_ can range from 2 to *k*. With the summing over all such groups explicitly shown, Eq. (64) becomes

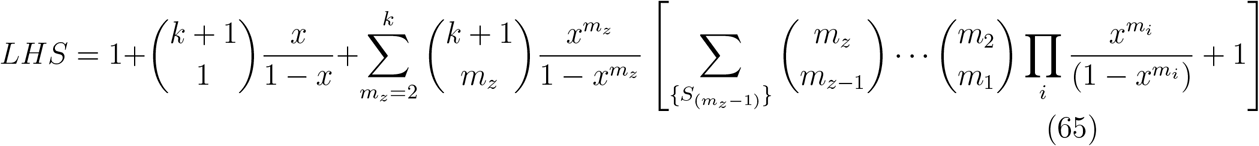

Note that 1 is added manually inside the square bracket to take care of the subsets with only one element *m*_*z*_, as these subsets are not taken into account by the summation term over the subsets {*S*_(*m*_*z*_*−*1)_}. The expression inside the square bracket is equal to the *LHS* of Eq. (49) with *n* = *m*_*z*_. Note that as positive integer *m*_*z*_ ≤ *k*, by assumption the identity Eq. (49) is true. So, substituting the form of *RHS* from Eq. (49) into the square bracket of Eq. (65), we have

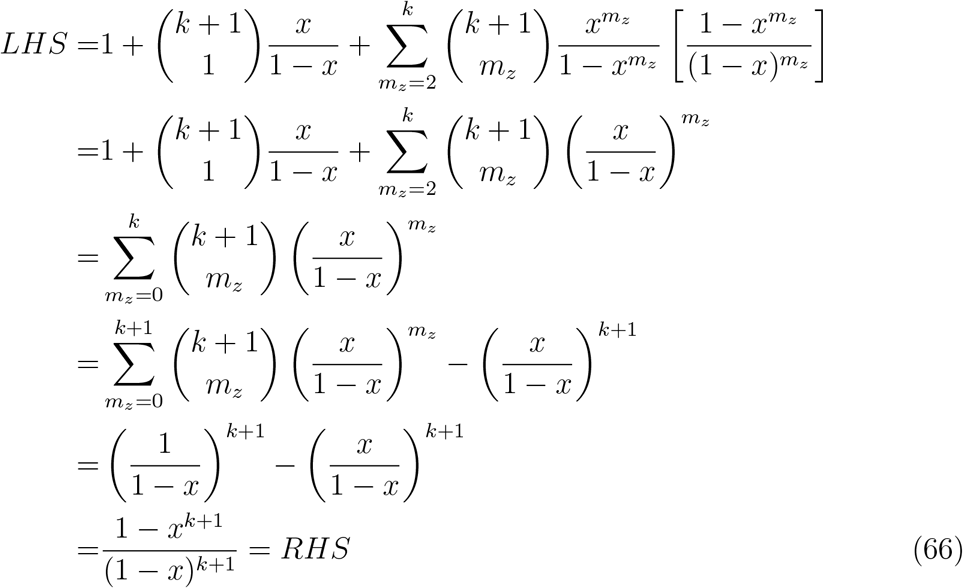

Hence, we have shown above that the identity is true for *n* = *k* + 1. Since integer *k* ≥ 2 is arbitrary, we have proved the identity in Eq. (49) using the method of strong induction.

## 7 Proof of the identity in Eq. (60)

Here is the proof of the identity using the method of mathematical induction:

For *n* = 2, {*S*}, the set of all the subsets of {1}, is simply {{1}}. Hence, the LHS of Eq. (60) will be equal to

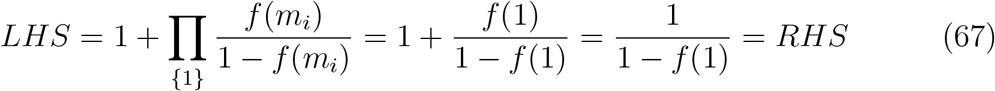

Let us assume that the identity is true for an arbitrary value of *n* = *k*. For, *n* = *k* + 1, the set {*S*} of all the subsets of {1, 2, 3, …*k*} can be divided into three disjoint subsets:

1. the set of all the subsets of {1, 2, 3, …*k* − 1}.
2. the set of all the subsets in (a) with an extra element *k* added to each of the subsets.

(c) {{*k*}}

Clearly, the contribution of (a) in the summation term in the LHS of Eq. (60) will be equal to that for *n* = *k*, that is equal to 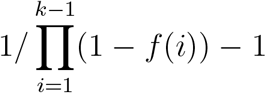. On the other hand, the contribution of (b) will be equal to that of (a) multiplied by an additional factor 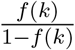. The contribution from (c) will be simply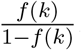. Adding all the three contributions,

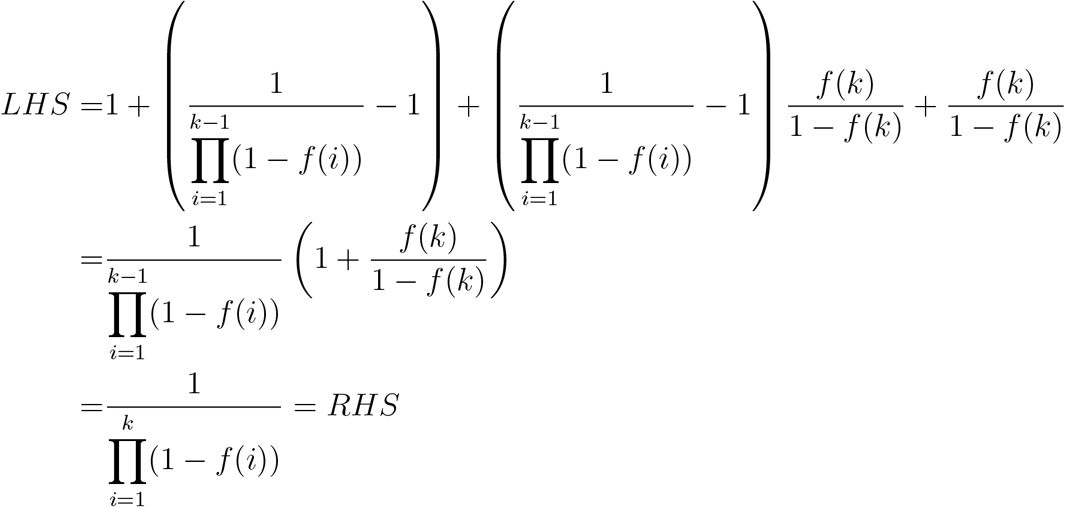

Hence, the identity is true for *n* = *k* +1. Thus from the principle of mathematical induction, the identity is proved for any integer *n >* 1.

## 8 Saturation of time dependent *p*_*r*_ and *k*, following calcium level saturation, in steady state_*t*_

**Figure 1:**
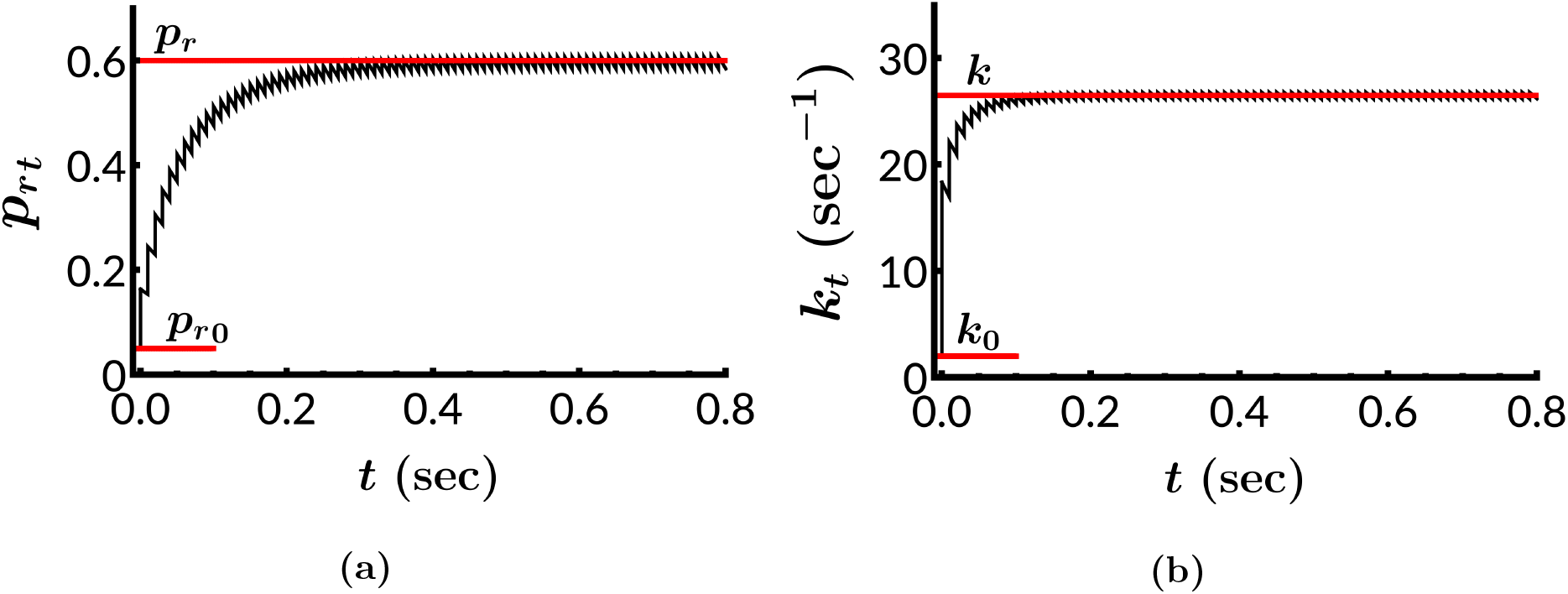
The time-dependence of *p*_*rt*_ and *k*_*t*_ are shown for the parameter values given in Table 2. of Ref. [1], for parallel fiber. The initial and steady state values are marked with red lines, and the corresponding values are mentioned.

The vesicle release probability *p*_*rt*_ and the rate of docking of vesicles *k*_*t*_ are dependent on the time *t*, through their dependence on time-varying Ca^2+^ level inside the presynaptic terminal. There are different models available in the literature to describe this period of plasticity. For the purpose of demonstrating the approach to steady state, we use one such model studied in the paper by Dittman and colleagues [1]. Two different types of calcium binding targets are considered, namely *F* and *D*. The *p*_*r*_ is related to the number *CaX*_*F*_ of Ca^2+^-bound-*F* molecules as 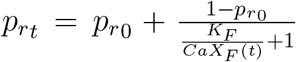. On the other hand, *k*_*t*_ is related to the number *CaX*_*D*_ of Ca^2+^-bound-*D* sites as 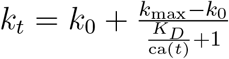. Given an AP train, with *m*^*th*^ AP arriving at *t* = *t*_*m*_, the dynamics of *CaX*_*F*_ and *CaX*_*D*_ are given by

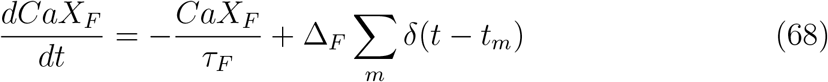

and,

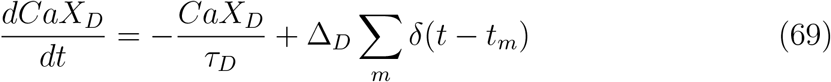

respectively. *CaX*_*F*_ (or *CaX*_*D*_) jumps by Δ_*F*_ (or Δ_*D*_) after each AP, and falls exponentially between two APs, with a decay constant *τ*_*F*_ (or *τ*_*D*_). In Fig. 1, for the parameter values given in Ref. [1] for parallel fiber, we have plotted the *p*_*rt*_ and *k*_*t*_ as a function of *t*. Note that the initial values at *t* → 0 are *p*_*r*0_ and *k*_0_ respectively. We see that *p*_*rt*_ → *p*_*r*_ and *k*_*t*_ → *k* in the long time limit. This steady state limit is the focus of the entire analysis described in this paper, in which the release probability and replenishment rate has attained the steady values *p*_*r*_ and *k* respectively.

## 9 Details of the experiments performed

We performed electrophysiological experiments in acute brainstem slices of juvenile mice (postnatal day 11) at MNTB-LSO synapses. These fast auditory synapses transfer inhibitory glycinergic information and can maintain reliable neurotransmission even during prolonged high-frequency stimulation [2, 3]. We refer the reader to the Methods section of Ref. [3] for details on the slice preparation and details of whole-cell patch-clamp recordings. The stimulation protocol consisted of repetitive tetanic AP bursts (100 Hz, 1 s) that were given 20 times. A 30-s rest period was introduced between repeats to allow for recovery from short-term depression. The number of fused SVs was determined as the ratio of the peak amplitude of evoked inhibitory postsynaptic currents (eIPSC) to the current produced by a single quantum(q) [pA/(pA/SV)]. Detailed information is available in Refs. [3, 4]. .

## 10 Kinetic Monte Carlo (KMC) code

The code used to simulate the distribution of quantal content for constant ISIs as well as Poisson train is as follows.

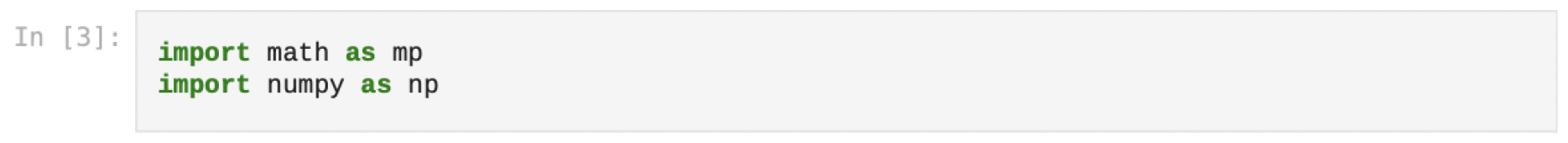

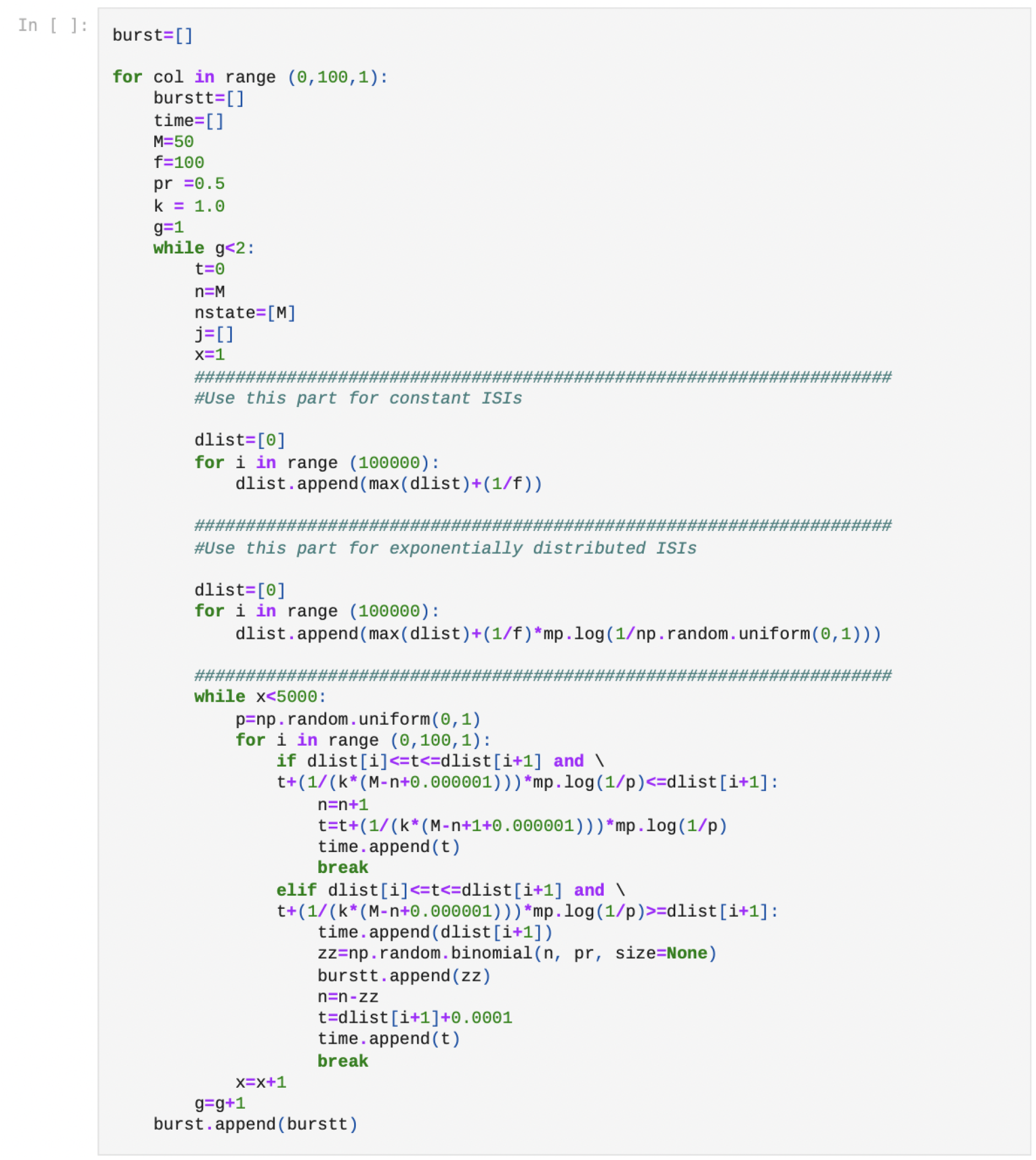

